# An integrated, scaled approach to resolve TSC2 variants of uncertain significance

**DOI:** 10.64898/2026.01.16.699909

**Authors:** Carina G Biar, Ziyu R Wang, Nathan D. Camp, Daniel L Holmes, Melinda K Wheelock, Sriram Pendyala, Abby V McGee, Pankhuri Gupta, Abbye E McEwen, Malvika Tejura, Marcy E. Richardson, Jamie D Weyandt, Taylor Coleman, Ross Stewart, Daniel Zeiberg, Allyssa J Vandi, Samantha Dawson, Predrag Radivojac, Lea M. Starita, Gemma L Carvill, Richard G. James, Douglas M Fowler, Jeffrey D Calhoun

**Author notes:** Corresponding authors: Jeffrey D Calhoun, Douglas M Fowler, Richard G. James. indicates equal contribution (Carina G Biar, Ziyu R Wang, Nathan D. Camp, Richard G. James, Douglas M Fowler, Jeffrey D Calhoun).

## Abstract

Obtaining a precise genetic tuberous sclerosis diagnosis is a challenge as many missense *TSC2* variants are variants of uncertain significance (VUS). VUS in *TSC2* have been resolved by one-at-a-time functional assays, but these assays cannot scale to the 3,634 *TSC2* missense VUS observed so far. To address this challenge, we used massively parallel sequencing to measure the steady-state abundance of almost 9,000 *TSC2* missense variants and developed an mTOR pathway activity assay using genome editing and cell sorting to generate activity scores for 391 missense variants. 1,288 of 8,891 (14.49%) missense variants assayed had altered TSC2 abundance, and 69 of 391 (17.65%) missense variants assayed had altered mTOR pathway activity. Calibration and integration of these data into classification of variants identified in a clinical cohort putatively reclassified 212 of 276 (76.8%) *TSC2* missense VUS. These datasets will lead to improved genetic diagnosis of tuberous sclerosis with potential positive impacts on the clinical management of patients and their families.

## Introduction

Tuberous sclerosis, a genetic disorder affecting multiple organ systems characterized by epilepsy and benign tumors, is caused by pathogenic variants in two mTOR pathway genes, *TSC1* and *TSC2*.^1,2^ *TSC1* and *TSC2* are also listed on the American College of Medical Genetics (ACMG) secondary finding list, as driver mutations in these genes are associated with increased risk for developing certain cancers such as early-onset renal cell carcinoma.^3^

Genetic diagnosis plays a major role in confirmation of a clinical diagnosis of tuberous sclerosis which can be challenging due to phenotype variability (**Figure 1A**).^4,5^ Genetic diagnosis of tuberous sclerosis increases access to precision treatments such as rapamycin analogs which inhibit mTOR.^6^ Of the 5,011 missense variants of *TSC2* in ClinVar, 174 (3.47%) are pathogenic or likely pathogenic (PLP), 223 (4.45%) are benign or likely benign (BLB), but 4,614 (92.08%) are either variants of uncertain significance (VUS) or have conflicting classifications. VUS arise when there is insufficient evidence to classify a variant as either pathogenic or benign or, more rarely, when there is conflicting evidence. Some *TSC2* VUS have been resolved by adding evidence from low-throughput functional studies in cell-based assays.^7–13^ This important work describing how *TSC2* missense variants impact mTORC1 activity and TSC2 protein abundance in cell-based assays is the current gold standard and has been used for many years to inform variant classification. However, the rapid accumulation of VUS is incompatible with the throughput of these classical cell-based assays. Meanwhile, multiplexed assays of variant effect (MAVEs), which harness high-throughput DNA sequencing to functionally characterize thousands of variants simultaneously, offer a tangible solution.^14^

**Figure 1.**
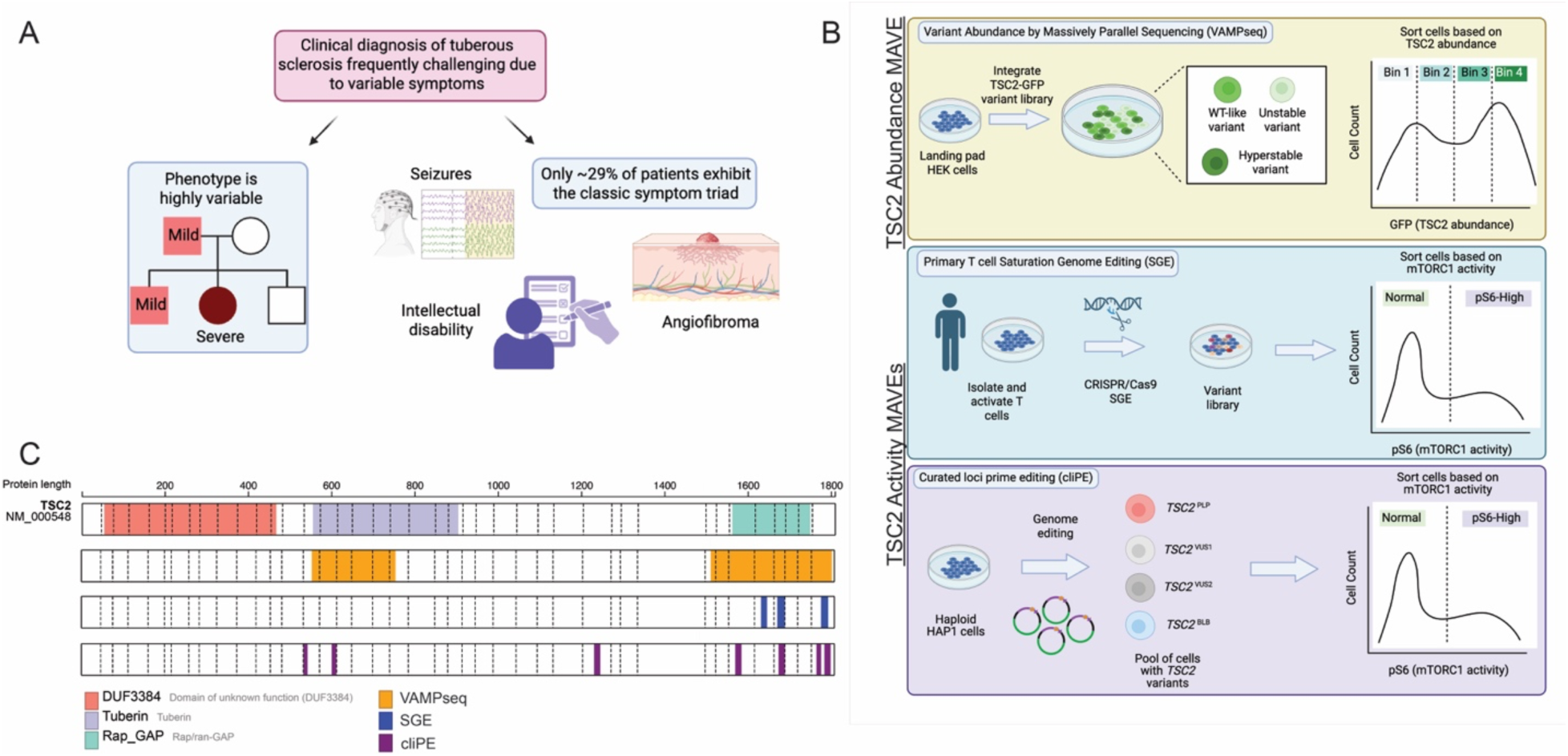
Prioritization of tuberous sclerosis related gene *TSC2* for generation of multiplexed assays of variant effect. (**A**) Major factors influencing prioritization of *TSC2* for MAVE development. (**B**) Introduction of three TSC2 MAVEs. Variant effect data for TSC2 abundance were generated using a VAMP-seq workflow. Two independent assays of TSC2 activity were performed in both primary donor T cells as well as HAP1s. (**C**) Diagrams demonstrating known functional domains of TSC2 as well as the regions targeted by the TSC2 MAVEs reported herein.

We developed two complementary MAVE-based approaches to resolve *TSC2* VUS (Figure 1B-C; Table 1). Some pathogenic TSC2 missense variants exhibit reduced protein abundance in heterologous systems, making abundance a promising molecular phenotype for identifying benign and pathogenic variants.^7–10,13^ Additionally, TSC2 functions as a negative regulator of mTORC1 signaling, making phosphorylation of the mTORC1 effector S6 (pS6) a direct readout of TSC2 activity (Figure S20). To measure TSC2 protein abundance, we employed variant abundance by massively parallel sequencing (VAMP-seq), which measures steady-state protein abundance by fusing variants to GFP followed by fluorescence-activated cell sorting (FACS) and sequencing.^15^ To assess TSC2 activity, we used endogenous genome editing combined with FACS for pS6, similar to our previous work on another mTOR pathway gene, *SZT2*.^16^ The editing approach overcomes a key limitation of VAMP-seq: its inability to capture splicing variants that can only be assayed at an endogenous locus. Given the variable phenotypic presentation of tuberous sclerosis even among individuals with identical variants, we implemented these TSC2 activity assays in two cell types. In primary T cells isolated from healthy donors, we used a cloning-free saturation genome editing (SGE) approach.^17^ In haploid HAP1 cells, we employed curated loci prime editing (cliPE), which uses prime editing to target specific regions enriched for VUS.^18^

**Table 1.** Overview of three TSC2 MAVE datasets.

|  | VAMP-seq | SGE | cliPE |
| --- | --- | --- | --- |
| Library type | cDNA | Genome editing (CRISPR/Cas9) | Genome editing (prime editing) |
| Coverage | Saturation | Saturation | Targeted |
| TSC2 variants (Total missense) | 8,891 | 268 | 153 |
| Selection | TSC2 abundance | TSC2 activity (mTORC1 activity via pS6 FACS) | TSC2 activity (mTORC1 activity via pS6 FACS) |
| Cell line/type | Landing pad HEK 293T | Primary donor T cells | HAP1 |

We measured the effect of 9,253 missense, synonymous, and nonsense variants on abundance using VAMP-seq and 522 variants on pS6 levels using SGE and cliPE. 1,288 missense variants reduced TSC2 abundance and 69 missense variants reduced TSC2 activity. Of these 69 variants, 34 altered both TSC2 abundance and mTORC1 activity. However, 31 variants reduced TSC2 activity but not abundance, and these variants clustered in the active site of the RAP-GAP domain. To reassess *TSC2* VUS, we developed a decision tree-based model to combine data from the assays. This model makes an initial assignment based on TSC2 abundance. Variants with an initial classification of functionally normal or indeterminate are replaced if SGE or cliPE data for that variant exists. OddsPath calibration is used to assign evidence strength, which varies from moderate evidence of benignity to strong evidence of pathogenicity depending on which assay is used for the final classification within the decision tree. We tested our decision tree method using 340 variants from a clinical cohort derived from a large commercial genetic diagnostic laboratory with an ACMG/AMP variant classification framework. 199 (72.1%) variants were putatively classified as likely benign, 13 (4.7%) as pathogenic/likely pathogenic, and 64 (23.1%) remained VUS. PS3/BS3 functional evidence codes derived from TSC2 MAVE data contributed to 91% (193/212) of classifications. Herein, we report the results of these three TSC2 MAVE datasets and discuss how these data will facilitate the resolution of missense *TSC2* VUS.

## Materials and Methods

### TSC2 VAMP-seq

#### Barcoding attB_IRES-mCherry plasmid

The VAMP-seq vector was modified based on a previously described system.^19^ Briefly, an 18-nucleotide randomized sequence, flanked by BsmBI recognition sites and fragment-specific overhangs, was used as the barcode insert. This fragment was assembled into the plasmid backbone, containing the *attB*-IRES-mCherry cassette and a C-terminally GFP-tagged construct, via complementary overhangs with 1:7 vector:insert ratio. The resulting product was transformed by two electrocompetent NEB 10-beta E.coli cells (NEB C3020) reactions recovered in 2ml SOC Media for one hour at 37C. Recovered cultures were plated to a 245mm LB+ AMP agar tray (Corning 07-200-600) and incubated at 30C overnight. Plasmid DNA was then isolated by maxipreparation (Zymo Research D4203). Barcode diversity was estimated by colony counting, yielding 2.73e9 unique barcodes for subsequent library cloning.

### Library cloning

Two site-saturation variant libraries were generated by Twist Biosciences based on the TSC2 open reading frame (NM_000548.5), C-terminally tagged with GFP. At each targeted site, we programmed a synonymous mutation, 19 missense mutations, a nonsense mutation, and a three base pair deletion. Library 1 is a site-saturation mutagenesis of amino acid residues 554-754 (187 sites passed QC) and library 2 is a site-saturation mutagenesis of amino acid residues 1512-1802 (286 sites passed QC). These two libraries were cloned to the above-mentioned barcoded plasmid using SapI golden gate cloning (NEB R0569S) followed by transformation by electroporation to achieve a barcode to variant coverage of 24x and 21x, respectively, for the two libraries. To associate barcodes with variants, we used PacBio SMRTbell sequencing.

Libraries were prepared according to the manufacturer’s protocol “Procedure & Checklist - Preparing SMRTbell® Libraries using PacBio® Barcoded Universal Primers for Multiplexing Amplicons.” This resulted in 930,059 reads for Library 1 and 244,393 reads for Library 2.

Barcodes were mapped to variants using Pacybara and only barcodes associated with a single unique variant at least two thirds of the time were used for VAMP-seq assays.^20^

### Tissue culture experiments and FACS sorting

Landing pad HEK 293T (HEK293T-LLP-iCasp9) cells were transfected using FuGene 6 with each barcoded variant library to achieve an estimated 20 cells per barcode coverage.^21^ At 48 hrs post-transfection, cells with successful integration were induced with 2 μg/mL doxycycline in media (DMEM + 10% FBS + 1% penicillin-streptomycin). After an additional 24 hours, cells were treated with 10 μΜ of AP1903/Rimiducid (MedChemExpress) to select against unrecombined cells. At day 4 post-transfection, cells were sorted on a BD FACSAriaII using an 85 μm or 100 μm nozzle into four bins (Bin 1: 0-25%; Bin 2: 25-50%; Bin 3: 50-75%; Bin 4: 75-100%) based on GFP:mCherry fluorescence via FACS to achieve at least two million cells per bin.^22^ Following sorting, the separate bins were grown out for several days to confluence in media (+ Dox) prior to harvesting genomic DNA from cell pellets (300g for 5 minutes).

### Genomic DNA isolation barcode sequencing

Genomic DNA was extracted from cell pellets of each bin using the Qiagen DNeasy Blood & Tissue Kit (Qiagen #695060) following the recommended protocol for purification of total DNA from animal blood or cells (Spin-column protocol using a maximum of five million cells per column), with the addition of 2 ul RNase A (20mg/ml) in the lysis step.

Two technical PCR replicates were performed following previously described protocols on each bin for each library to assess barcode correlation from amplification and sequencing.^22,23^

Each indexed sample was sequenced for barcodes using custom read and indexing sequencing primers NP0494, NP0495; NP0497, and NP0550 using a NextSeq 1000/2000 P3 50 cycles kit (Illumina).

### VAMP-seq data analysis and scoring

Variants were scored using previously published methods.^15^ Briefly, fastq files were generated using bcl2fastq v2.20 and paired reads were merged using PEAR v0.9.11. Fastq reads were mapped to barcodes from the Pacybara output and collapsed to generate count tables where each row represents a variant and each column is a single bin in a replicate experiment. Variant frequencies within each bin were calculated by dividing the count for that variant by the sum of all counts in that bin. Bin frequency values were used to generate weighted abundance scores for each variant by taking the frequency in bin 1 and multiplying by 0.25, bin 2 by 0.5, bin 3 by 0.75, and bin 4 by 1 and summing those values before dividing by the unweighted sum for that variant across all bins. The resulting values were min-max normalized so that WT is 1 and the median of the lowest 5% of missense values is 0 to generate the reported abundance scores. All VAMP-seq scores are reported in Table S1.

### Primary T cell saturation genome editing

#### CD4^+^ T cell isolation, activation, and culture conditions

Deidentified human peripheral blood mononuclear cells were acquired from healthy donors under informed consent from the Fred Hutch Specimen Processing and Research Cell Bank (protocol #3942). CD4^+^ T cells were isolated (Fred Hutchinson Cancer Research Center) using the EasySep Human CD4^+^ T cell enrichment kit (Stem Cell Technologies 19052). Cells were cultured at a density of 0.5e6 cells/mL in RPMI (ThermoFisher) supplemented with 20% FBS (Sigma), 1x Glutamax (ThermoFisher), 10 mM HEPES (ThermoFisher), 0.1% BME (ThermoFisher) and 50 ng/ml IL-2 (PeproTech). For activation, Dynabeads Human T-Expander CD3/28 (ThermoFisher) were added at 30 uL/1e6 cells. Beads were removed after 72 hours, and cells were allowed to rest and proliferate for 16-20 hours before transfection.

### Transfection of CD4^+^ T cells

100 uM crRNA (IDT) was complexed with 100 uM trRNA (IDT) at a ratio of 1:1 and incubated at 95C for 5 minutes, then allowed to cool at room temperature for 30-60 minutes. Recombinant Cas9 (UC Berkeley) was added at a guide:Cas9 ratio of 4:1 and the mixture was allowed to rest at room temperature for 15 minutes. oPool variant libraries (IDT) were designed to have two phosphorothioate bonds between the three terminal bases and were composed of all possible variants within an SGE region (Table S10). oPool libraries were added at a final concentration of 40 pmol/1e6 cells. CD4+ cells were resuspended in Maxcyte buffer (Maxcyte) and added to the Cas9/guide/library mix. This sample was then transferred to a Maxcyte assembly and transfected with the Expanded T Cell-1 protocol on a Maxcyte ExPERT GTx electroporator.

### Cell staining and sorting

Following transfection, cells were cultured for 6 days with media replacement every 48 hours. Approximately 1e6 cells were isolated for downstream genomic DNA and the remaining cells were fixed, permeabilized, and stained for phosphorylated S6 (Cell Signaling Technology 5316S). Stained cells were analyzed on a BD FACSAria II flow cytometer and cells were sorted on the highest and lowest 25% of phospho-S6 intensity.

### Genomic DNA isolation

Genomic DNA was isolated from presorted cells using the Quick-DNA miniprep kit (Zymo Research). For fixed cells, approximately 1-2e6 cell pellets were suspended in 250 uL of PBS containing 200 ug of RNAse A (Sigma) and 200 ug Proteinase K (Zymo Research). Cells were incubated at 37C for 30 minutes. 250 uL of 2x lysis buffer containing 60 mM Tris pH 8, 20 EDTA, and 2% SDS was added and cells were incubated overnight at 65C with shaking at 800 rpm. 250 uL of ethanol was then added to the sample and genomic DNA was isolated with the Quick-DNA miniprep kit.

### Illumina sequencing

Each SGE region was first amplified with intron-specific primers amplifying the entire exonic region with NEBNext Ultra II Q5 high fidelity DNA polymerase (New England Biolabs) (Table S9). This sample was then purified with Agencourt AMPure XP reagent (Beckman Coulter) at a 0.9x beads to sample ratio. These samples were then amplified with locus-specific primers containing Illumina adapter sequences. These samples were purified with 0.9x AMPure beads and reamplified with primers containing Illumina p5 and p7 sequences and a unique sample index. These samples were purified one final time with 0.9x AMPure beads. The amplicon library was verified for purity by PAGE gel, quantified by Qubit 3 (ThermoFisher), pooled, and paired-end sequenced on an Illumina NextSeq 2000 with the XLEAP P2 300 cycle kit (Illumina).

### Data analysis and scoring

After Illumina sequencing, individual replicates (n=4 replicates: 2 donors, 2 replicates each donor) were de-multiplexed and Illumina reads were merged with the PEAR software package. We then used custom Python scripts to isolate and count variants in the paired-end reads for the pS6-Hi and pS6-Lo populations. We then used the CountESS software package to generate functional scores. Briefly, we normalized variant counts within each sample, then calculated the replicate score as the ratio of variants in the pS6-Hi population compared to that in the pS6-Lo population. Scores were then scaled to the bottom 3 (min) and top 3 (max) scores of that region, then scaled again to the synonymous variant within that region. This score was then log transformed and the final pS6 score was calculated as the mean of these scores across all 4 replicates. All SGE scores are reported in Tables S1 and S2.

### Curated loci prime editing

cliPE is a modification of a published method for saturation prime editing.^24^ For a detailed cliPE protocol, please see Biar *et al.* or the supplementary methods.^18^

### OddsPath calibration

Scores for each MAVE were calibrated using standard methods.^25^ For VAMP-seq, we used a threshold of <0.25 to determine variants with abnormal, or low, abundance. This stringent threshold was selected to maximize separation of ClinVar BLB and PLP missense variants. We assigned variants scoring between 0.25 and 0.75 for abundance as indeterminate. Variants with abundance scores >0.75 are considered as having normal abundance, coinciding with the lower 95% confidence interval for synonymous variants. For SGE and cliPE, we developed single thresholds to classify variants as either having normal or abnormal activity. For SGE, variants with TSC2 activity scores <2 were classified as having normal activity, while variants with scores >2 were classified as having abnormal activity. For cliPE, variants with TSC2 activity scores <0.25 were classified as having normal activity, while variants with scores >0.25 were classified as having abnormal activity. For all assays, OddsPath calculations were performed using the current ClinGen-approved calibration framework (Table 2).

**Table 2.** Odds of Pathogenicity (OddsPath) calculations for individual MAVE datasets. *One ClinVar BLB variant, p.S641I, was excluded from these calculations; please see Discussion for rationale.

|  | ClinVar BLB | ClinVar PLP | OddsPath_PS3 | OddsPath_BS3 |
| --- | --- | --- | --- | --- |
| VAMP-seq | 54* | 80 | 33.1 ('strong') | 0.343 ('supporting') |
| SGE | 6 | 16 | 5.6 ('moderate') | 0.094 ('moderate') |
| cliPE | 6 | 10 | 4.5 ('moderate') | 0.113 ('moderate') |

### Variant classification on Ambry cohort

Preliminary Ambry classifications for *TSC2* missense variants that have been observed in individuals tested at Ambry genetics (n=340) were performed utilizing the Ambry Classifi framework based on ACMG/AMP guidelines and implementing a points-based approach.^26,27^ Diagnostic criteria for application of phenotype-based evidence was based on international consensus criteria.^28^ Population-based evidence thresholds were established within Ambry’s Classifi program using gene-specific Whiffin-Ware calculations.^29^ For the final classifications, *in silico* prediction-based points were assigned based on AlphaMissense thresholds that were calibrated for *TSC2* and functional weight was incorporated using the calibrated points-based approach described in **Table 2** (range of -2 to 4).^25,26^

### Variant classification on all other missense variants assayed

A simplified version of the points-based system used for the Ambry cohort was used for variants that had not been previously observed in the Ambry cohort. Where available, a calibrated *in silico* predictor optimized for *TSC2* was generated from AlphaMissense scores.^30^ Based on the strength of evidence calculated for the functional datasets (**Table 2**), variants were assigned a range of points between -2 and +4.^25^

## Results

### Development of multiplexed assays of variant effect for TSC2

Pilot VAMP-seq experiments showed WT TSC2-GFP expressing cells were separable from cells expressing a premature termination codon (PTC) variant (**Figure S21-22**). Cells expressing a pilot library of 47 variants were distributed broadly with a peak corresponding to cells with low TSC2-GFP abundance (**Figure S23**). We sorted cells into bins based on GFP fluorescence and performed amplicon sequencing to confirm that WT expressing cells were enriched in high GFP bins while PTC variants were enriched in low GFP bins (**Figure S24**). Based on these results, we proceeded to perform VAMP-seq at saturation scale. Synonymous TSC2 variants (n=362) were distributed with a relatively narrow range of scores (mean=0.998; standard deviation=0.096) (**Figure 2A**). In contrast, the scores for missense variants (n=8,891) were bi-modally distributed (mean=0.751; standard deviation=0.347), with one mode overlapping synonymous variants. Some missense variants had intermediate abundance scores, suggesting that they may be hypomorphic in terms of protein abundance. 1,288 of 8,891 (14.49%) variants had low abundance in the VAMP-seq assay. 5,740 (64.56%) variants had normal abundance; the remaining 1,863 (20.95%) variants were considered indeterminate. 27 of 8,891 missense variants may have increased protein abundance (TSC2 abundance >1.35, exceeding the right 95% confidence interval for synonymous variants). However, it is unclear if these variants impact TSC2 activity. Only one variant (p.Gly1642Asp) among these is known to be pathogenic and had low activity in the SGE assay. VAMP-seq abundance measurements weakly correlated (Pearson correlation coefficients between -0.18 and -0.23) with evolutionary conservation metrics and moderately correlated with variant effect predictor (VEP) scores (Pearson correlation coefficients ranging between -0.29 and -0.54); AlphaMissense showed the strongest negative correlation among VEPs (**Figure 3**).^31^

**Figure 2.**
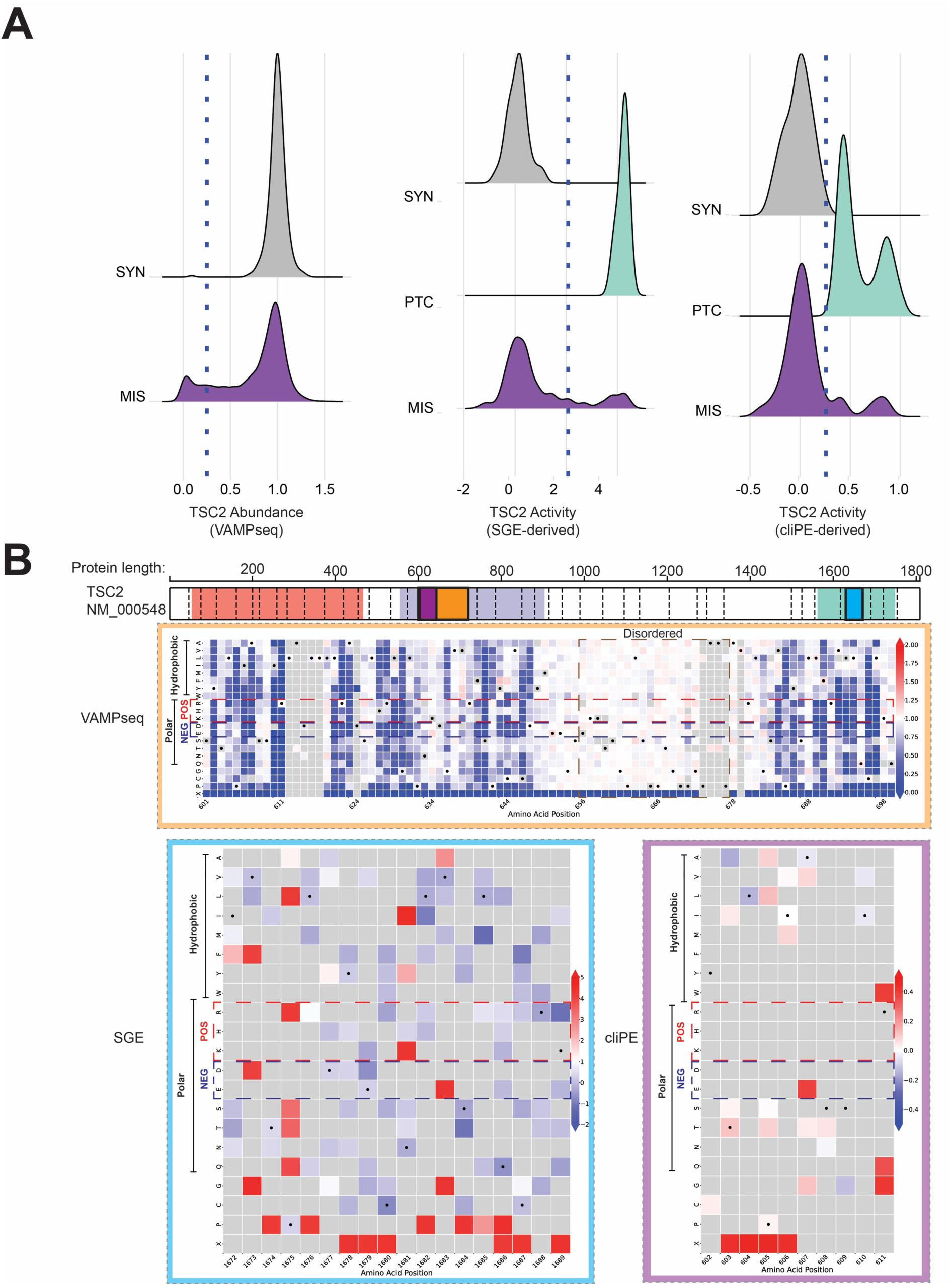
TSC2 variant abundance and activity mapping. (**A**) Distributions of variant effect scores by variant class. On the left, a blue dotted line shows the threshold for low abundance variants. In the middle and right panels, a blue dotted line shows the threshold separating variants with normal activity from those with abnormal activity. SYN=synonymous. PTC=premature termination codon. MIS=missense. (**B**) Example heatmaps showing individual variant effect scores. Boxes overlaid on the TSC2 diagram denote which regions are displayed. Amino acid residues are grouped by biochemical properties. Positively charged residues are annotated with a red box with dotted lines and negatively charged residues are annotated with a blue box with dotted lines. A disordered region is annotated with a brown box with dotted lines; substitutions within and adjacent to this disordered region have little to no effect on TSC2 abundance.

**Figure 3.**
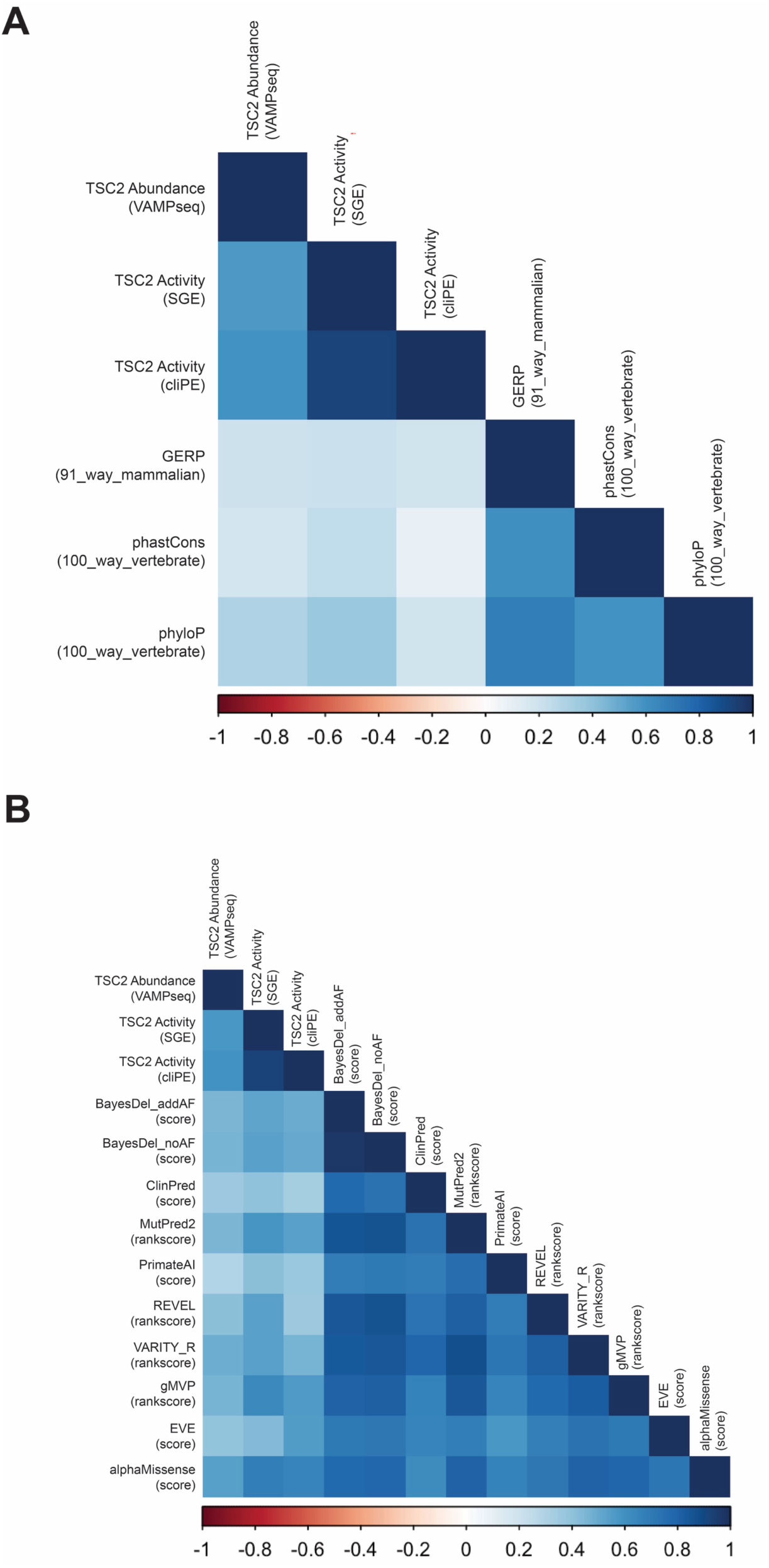
Correlation of TSC2 MAVEs with evolutionary conservation metrics and variant effect predictors. (**A**) Correlation plot of TSC2 MAVE datasets and evolutionary conservation metrics. (**B**) Correlation plot of TSC2 MAVE datasets and variant effect predictors. The inverse of the TSC2 abundance VAMP-seq score was used to calculate correlation coefficients in the first column.

As a proof-of-concept for a TSC2 activity MAVE, we first performed pilot experiments with a single guide RNA targeting TSC2 exon 17. We demonstrated that cells with NHEJ-induced indels had higher levels of pS6 relative to untransfected cells (**Figure S26-27**). Based on this, we next generated libraries of variants using two methods in both primary T cells and HAP1 cells to capture potential cell-type specific variant effects. We generated TSC2 activity scores for 522 variants, including 391 missense variants, 64 synonymous variants, and 67 PTC variants.

The activity scores of synonymous and PTC variants comprised distinct populations with non-overlapping distributions, showing these assays distinguished functionally normal from functionally abnormal variants (**Figure 2A**).The activity score distributions for all missense variants were bimodal with one mode overlapping the synonymous variant distribution and the other overlapping the PTC variant distribution, revealing that some missense variants retain activity while others have abnormal activity. 52 of 268 (19.4%) missense variants exhibited abnormal activity in the SGE assay. 22 of 153 (14.38%) missense variants had abnormal activity in the cliPE assay. Activity scores were weakly to moderately correlated (Pearson correlation coefficients between 0.1 and 0.38) with measures of evolutionary conservation (**Figure 3**). In general, activity scores were positively correlated with VEP scores, with AlphaMissense having the strongest correlation (Pearson correlation coefficients between 0.38 to 0.68; AlphaMissense/SGE = 0.68, AlphaMissense/cliPE = 0.65). Overall, AlphaMissense correlates most strongly with our abundance, SGE-derived activity, and cliPE-derived activity datasets, suggesting it may be the optimal VEP to calibrate and use to assign *in silico* pathogenicity evidence in an ACMG/AMP variant classification framework.

### Approximately one-third of residues assayed for abundance were fully or partially intolerant to amino acid substitution

In total, 377 TSC2 residues were tolerant to amino acid substitution (<20% of substitutions assayed resulting in loss of abundance). We identified 48 residues which were highly intolerant to amino acid substitution (>60% of substitutions result in low protein abundance). The remaining residues (n=61) partially tolerate substitution (between 20 and 60% of substitutions result in low protein abundance). To better understand these variants with reduced abundance, we analyzed them in the context of the AlphaFold2 predicted structure for TSC2. Many tolerant residues mapped to predicted disordered regions, such as between residue 650 and 680 (**Figure 2B**). Similarly, many tolerant residues mapped to the C-terminal region spanning residues 1724-1805, which is mostly disordered. However, some predicted structured regions, such as the alpha helix spanning residues 704-716, contained multiple tolerant residues. Highly intolerant and partially tolerant residues mapped to structured regions including alpha helices (such as residues 582-598) and beta sheets (such as residues 1646-1651). Examples of residues highly intolerant and partially intolerant are 611 and 610, respectively. Residue 611 is a well-known residue in the tuberous sclerosis genetics field with 4 known ClinVar PLP variants (R611W, R611P, R611Q, and R611G).

### TSC2 variant abundance and activity are strongly correlated

We next analyzed the relationship between TSC2 variant abundance and activity as measured in primary T cells and HAP1 cells (**Figure 4**). Abundance was strongly correlated with activity in primary T cells (Pearson correlation = -0.57) and in HAP1 cells (Pearson correlation = -0.69, **Figure 4A-B**). The SGE and cliPE datasets correlate very strongly with a coefficient of 0.92, suggesting few if any cell-type-specific variant effects (**Figure S28**). Thus, the majority of variants have concordant effects on abundance and activity as well as on activity between cell types. 31 variants exhibited normal abundance yet reduced activity, and we hypothesized that these variants directly impacted TSC2 catalytic activity. Indeed, these were located near the active site of the RAP-GAP domain (**Figure 4A-B**).^32^ Only residues within an alpha helix, the beginning of the adjacent beta sheet, or the linker region between these two secondary structure elements resulted in reduced activity variants that retained normal abundance. Of these inactive yet abundant variants, all but one mapped to highly evolutionarily conserved residues.

**Figure 4.**
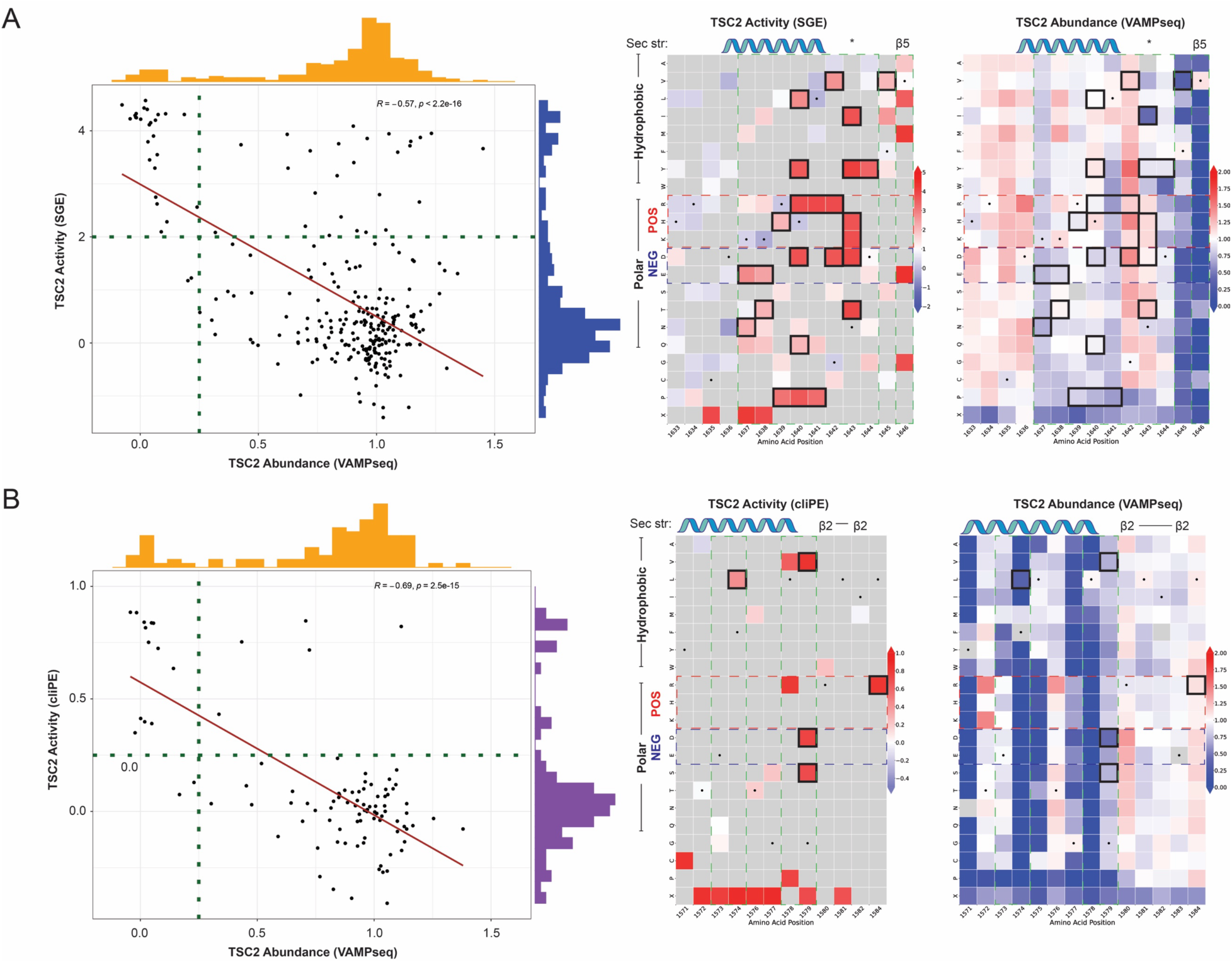
TSC2 abundance and activity are strongly correlated. (**A**) VAMP-seq and SGE comparison. Pairwise correlation of the VAMP-seq TSC2 abundance dataset with the SGE TSC2 activity dataset (left). Dotted lines indicate the assay thresholds separating variants with normal abundance or activity from those with altered abundance or activity. Example heatmaps spanning amino acid residues 1633 to 1646 of the RAP-GAP domain (right). Variants meeting criteria for normal abundance and altered activity are shown highlighted with black squares. Amino acid residues are grouped by biochemical properties. Positively charged residues are annotated with a red box with dotted lines and negatively charged residues are annotated with a blue box with dotted lines. Secondary structure is annotated above each heatmap. Asterisk designates the asparagine residue (p.N1643) which is a critical active site for RAP-GAP domain function. Regions of high evolutionary conservation are annotated with a green box with dotted lines; most substitutions within and adjacent to this region have little to no effect on TSC2 abundance. (**B**) VAMP-seq and cliPE comparison. Pairwise correlation of the VAMP-seq TSC2 abundance dataset with the cliPE TSC2 activity dataset (left). Example heatmaps spanning amino acid residues 1571 to 1584 of the RAP-GAP domain (right). Annotations of the plots are as described in (**A**).

### TSC2 variant effect data improve genetic diagnosis of tuberous sclerosis by resolving missense VUS

We evaluated the utility of functional data for preliminary reclassification of *TSC2* VUS using the ClinGen-approved Odds of Pathogenicity (OddsPath) framework. We analyzed the abundance and activity scores of *TSC2* missense BLB (n=54) and PLP (n=80) control variants found in ClinVar (**Figure 5**). For both abundance and activity scores, BLB and PLP controls minimally overlapped, with the exception of a proportion (31/80; 38.75%) of PLP variants with normal abundance. The separation of PLP and BLB variants by abundance and activity scores suggested that the data would have utility for variant classification and VUS resolution. Thus, we calculated the strength of evidence generated by each dataset using the OddsPath approach (**Table 2**).^25^ Under this framework, reduced abundance variants receive strong evidence of pathogenicity (OddsPath 33.1; PS3_strong) and normal abundance variants receive supporting evidence of benignity (OddsPath 0.343, BS3_supporting). Reduced activity variants in either the primary T cell or HAP1 activity assays received moderate evidence of pathogenicity (OddsPath 4.5,5.6; PS3_moderate) and normal activity variants in either assay received moderate evidence of benignity (OddsPath 0.113, 0.094; BS3_moderate).

**Figure 5.**
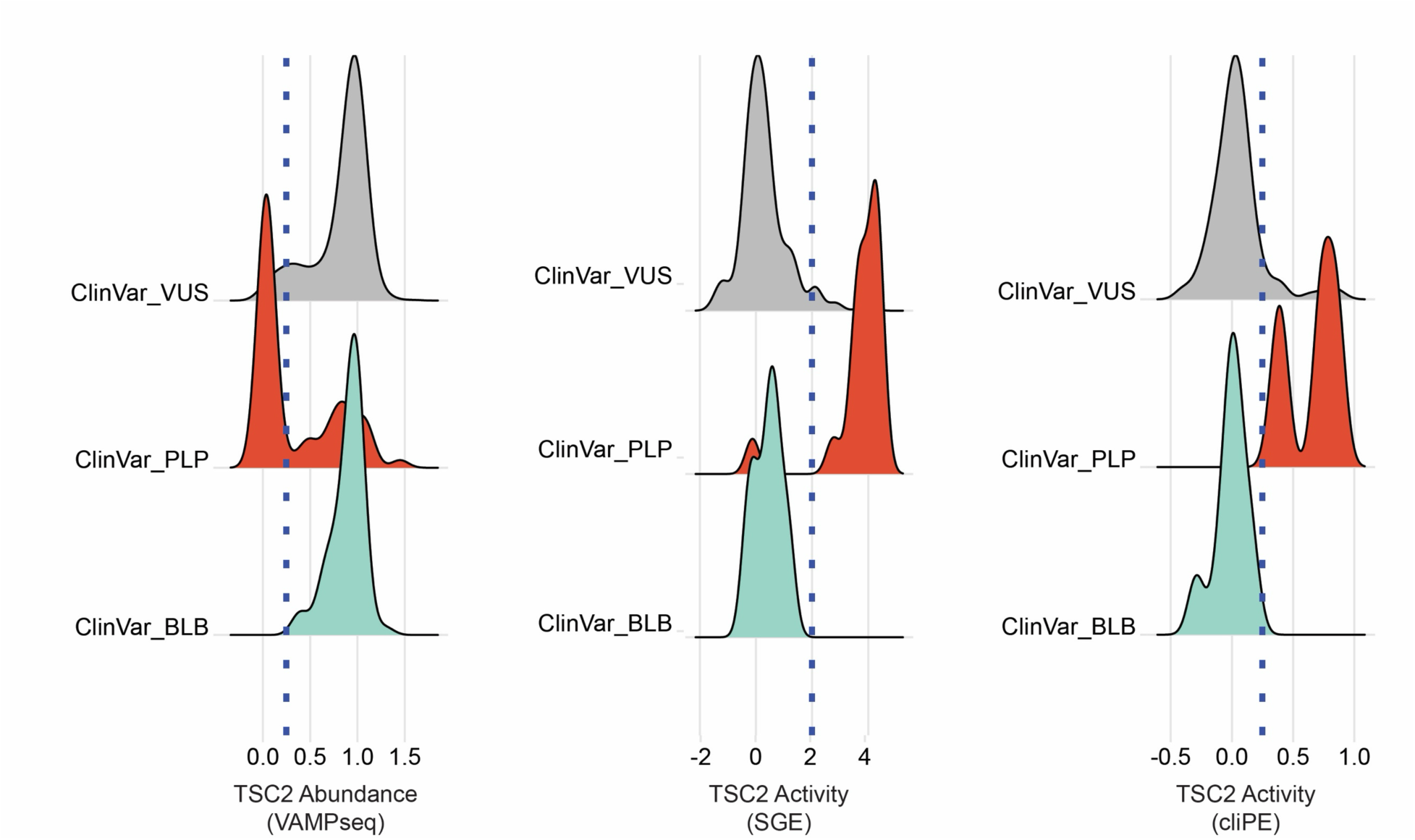
ClinVar variants of known pathogenicity or benignity have distinct distributions across TSC2 MAVE datasets. Distributions of variant effect scores by ClinVar annotation. On the left, a blue dotted line shows the threshold for low abundance variants. In the middle and right panels, a blue dotted line shows the threshold separating variants with normal activity and those with abnormal activity.

To combine evidence from the abundance and activity assays, we developed a decision tree to guide how the data we present can be used in the ACMG/AMP framework (**Figure 6A**). Using calibrated functional data together with Ambry’s evidence and computational predictions, 212 of 276 (76.8%) *TSC2* missense VUS with available functional data were putatively reclassified under ACMG/AMP (v3.0) criteria. Of these, 199 (72.1%) qualified as likely benign, 13 (4.7%) as pathogenic/likely pathogenic, and 64 (23.1%) remained VUS. Computational evidence contributed to 90.5% (192/212) reclassifications, and functional data contributed to 91% (193/212). Variants remaining VUS had indeterminate functional results (n=31), conflicting functional and computational evidence (n=23), or insufficient evidence for reclassification (n=10). The variant classification using this approach is shown in **Figure 6B**.

**Figure 6.**
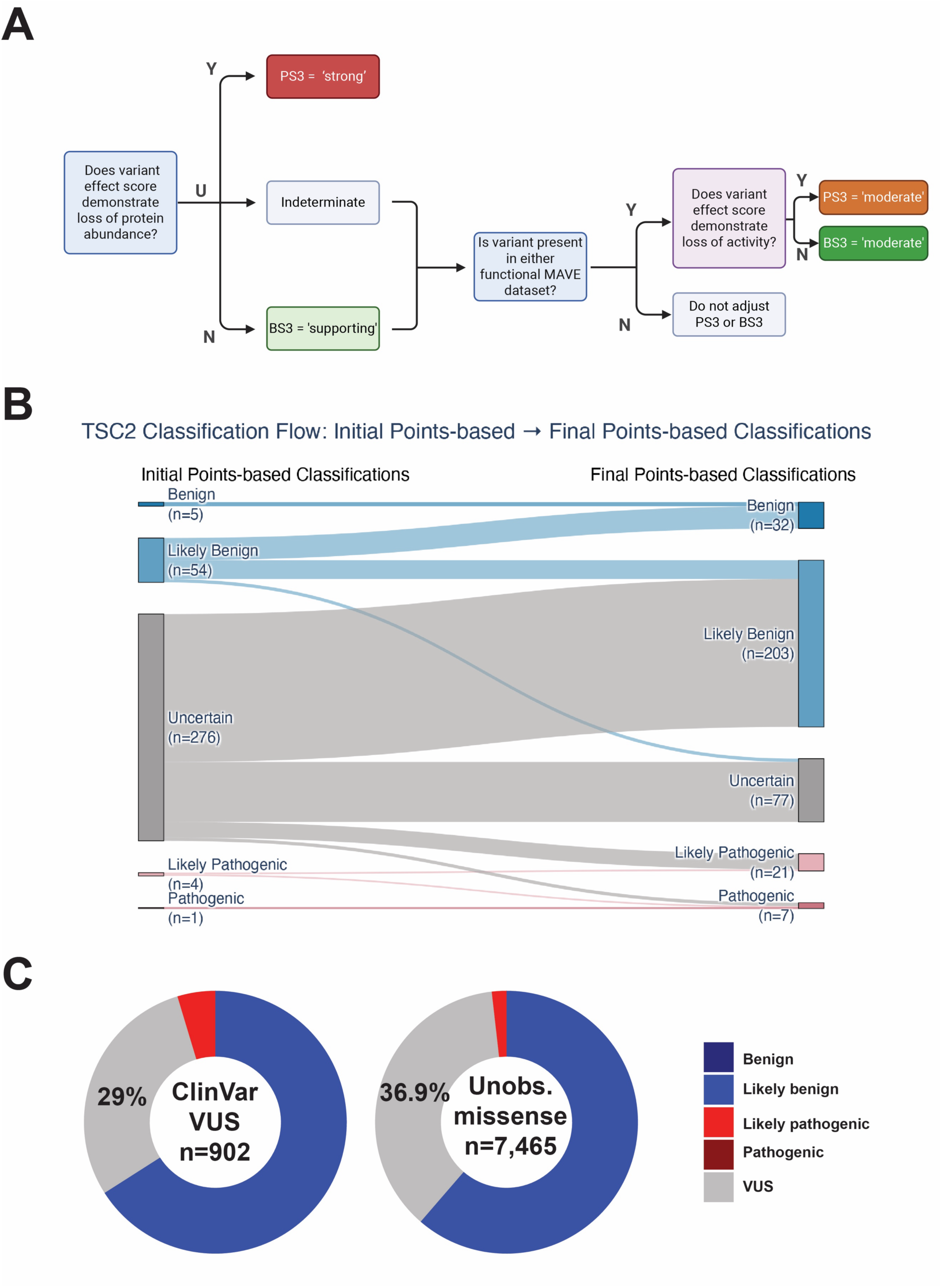
Employing *TSC2* MAVE data for resolution of missense VUS in a standard framework. (**A**) Proposed decision tree for utilization of MAVE datasets during standard variant classification. (**B**) Points-based implementation of ACMG/AMP (v3.0) variant classification for Ambry cohort. On the left are initial points-based classifications; on the right are final, putative points-based classifications using calibrated *in silico* VEP evidence and PS3/BS3 evidence derived from the decision tree in **A**. This framework enabled putative reclassifications of VUS to either likely pathogenic or likely benign, as well as up- and downgrading some likely benign and likely pathogenic variants. (**C**) Variant classification using *in silico* VEP prediction evidence and PS3/BS3 evidence codes derived from the decision tree in **A**. BS3/PS3 evidence codes enable reclassification of missense variants to either likely pathogenic or likely benign.

Many variants in our MAVE datasets were not in our clinical dataset. Experimental and predictive data were used to determine whether MAVE data could move these variants from VUS to other categories (Table S7 and **Figure 6C**). Of 902 missense ClinVar VUS with variant effect data, we putatively classified 595 (65.96%) variants as likely benign, 265 (29.38%) variants as uncertain, and 42 (4.66%) variants as likely pathogenic. Predictive data were available for 100% of ClinVar missense VUS. We similarly investigated unobserved variants, or variants which have yet to be observed in an individual in gnomAD or ClinVar. Of 7,465 unobserved missense variants, we assigned putative classifications of 4,579 (61.3%) variants as likely benign, 2,752 (36.9%) variants as uncertain, and 134 (1.8%) variants as likely pathogenic (**Figure 6C**). Predictive data were available for 1,357 (18.2%) unobserved missense variants.

## Discussion

We developed three multiplexed assays to resolve *TSC2* VUS. We used VAMP-seq to measure the abundance of 8,891 unique missense variants in an N-terminal portion of TSC2 and the full RAP-GAP domain. We used SGE and cliPE to measure the S6 phosphorylation, a proxy for the activity of TSC2, on 522 variants in primary T cells and HAP1 cells, respectively. Our data revealed that some variants in the RAP-GAP domain cause loss of TSC2 activity with no change in protein abundance. This suggests that TSC2 activity and abundance data are complementary, which argues that generating variant effect data for both abundance and activity is optimal for *TSC2*. In a clinical cohort, we reclassified more than 75% of VUS and variant effect data contributed to over 90% of the reclassifications, demonstrating the clinical utility of our MAVE datasets.

Our data can empower the precise genetic diagnosis of tuberous sclerosis by reducing VUS burden and by providing evidence to avoid future VUS. Reducing VUS in *TSC2* is clinically meaningful because PLP variant diagnosis can be used to prescribe mTOR inhibitor therapy, a first-line therapy for tuberous sclerosis.^33^ In addition, resolving VUS to PLP can increase involvement in clinical trials for new precision therapies and empower family planning. Resolving VUS to BLB can eliminate worry, prevent unneeded treatment, or suggest the need for further genetic testing and clinical workup. It is worth emphasizing that, from a clinical genetics standpoint, the greatest strength of the TSC2 abundance variant effect dataset is detecting pathogenic variants (specificity=98.18%). While the assay generates evidence towards benignity at supporting evidence strength (sensitivity=61.25%), appropriate care must be taken when using this benign evidence for variant classification.

In the course of our work, we identified some possible errors within ClinVar. First, p.S641I, labeled as a low abundance variant by VAMP-seq, was classified as a benign variant in ClinVar. p.S641I has been previously published as a pathogenic missense variant and further listed as either likely pathogenic or pathogenic in multiple unique entries within the *TSC2* LOVD database.^34,35^ Second, p.S540P, a variant with reduced TSC2 activity as measured in our cliPE assay, is listed as a benign variant in ClinVar. Lastly, p.Q1686K is listed as a likely pathogenic variant within ClinVar. We infer that the known pathogenic variant p.Q1686P likely contributed to this classification of p.Q1686K. However, in our primary T cell activity assay, p.Q1686P, but not p.Q1686K, met criteria for abnormal activity. We also establish that many residues in TSC2 are partially tolerant to amino acid substitution, suggesting PM5 evidence should be used with caution for TSC2.

Our work has several limitations. First, VAMP-seq is a cDNA-based assay and therefore is not appropriate for examining potential splice-altering variants. SGE and cliPE, by contrast, are based on endogenous editing and thus can capture the effect of splicing variants. Second, while both VAMP-seq and SGE are saturation approaches and therefore both prospective and retrospective, it is important to mention that cliPE is an inherently retrospective approach. In cliPE, we generate variant libraries based on clinically observed variants in databases such as ClinVar, LOVD, and gnomAD at one point in time. As more individuals are sequenced, we are likely to uncover novel variants in *TSC2*. Saturation approaches like VAMP-seq or SGE inherently provide data to help us understand these variants, whereas additional cliPE experiments would need to be performed to include such variants. We recommend cliPE as a good alternative to SGE when budget is a limiting factor; SGE should be considered if funding permits. Finally, low-passage HAP1 cells were used for cliPE, but we did not select for ploidy using flow cytometry or pharmacology.^36,37^ As *TSC2* is an autosomal dominant disorder, the ploidy of cells is unlikely to be a major confounding factor. However, we cannot rule out a small effect of ploidy on our assay results, particularly for putative partial loss-of-function variants or dominant negative variants. Our primary goal was developing a MAVE which could distinguish complete loss-of-function missense variants from variants retaining function. Future work will be necessary to develop assays that are calibrated to test for partial loss-of-function or dominant negative variants.

This study builds on the excellent work by the Nellist lab and others to functionally characterize individual missense variants in *TSC2*.^7^ We have made our variant effect data available in the supplementary information (Table S1). Our work is registered on the MAVE Registry and our data has been deposited in MAVEdb to share with the community.^38,39^ Future directions of this work include forming a Clingen Variant Curation Expert Panel to generate a custom adaptation of the ACMG/AMP guidelines optimized for *TSC2*. It will be important to complete additional VAMP-seq, SGE, and cliPE studies to generate a complete variant effect map for the full *TSC2* gene.

Pathogenic variants in at least 16 mTOR pathway genes result in mTORopathies, a group of neurodevelopmental disorders commonly associated with epilepsy, in both children and adults.^40^ We also view this study as a proof-of-concept towards comprehensive VUS resolution in the mTORopathies. The methods utilized herein are generalizable and could be used to study mTORopathy genes with hundreds of missense VUS such as *MTOR* and *PIK3CA*, perhaps increasing access to mTOR inhibitors and other forms of precise clinical management.

## Web resources

gnomAD v4 (https://gnomad.broadinstitute.org/); accessed 5 Jan 2024

Regeneron Genetics Center (RGC) Million Exome Variant Browser (https://rgc-research.regeneron.com/me/license-and-terms-of-use); accessed 5 Jan 2024

ClinVar (https://www.ncbi.nlm.nih.gov/clinvar/); accessed 24 Oct 2022 for initial list of BLB, PLP, and VUS for design of epegRNA libraries; accessed 18 Dec 2023 for final list of BLB, PLP, and VUS

*TSC2* LOVD gene-specific database (https://databases.lovd.nl/shared/genes/TSC2); accessed 8 Jan 2024

IGVF Portal (https://data.igvf.org/) MAVEdb (https://mavedb.org/)

Some figure panels were created with BioRender.com.

## Data Availability

VAMP-seq data is available via the IGVF Portal (Analysis set: IGVFDS5595BTYJ; Score file: IGVFFI9747KART). SGE data is available via the IGVF Portal (Analysis set: IGVFDS1782FCXW; Score file: IGVFFI3097DFGF). cliPE data is available at MAVEdb (urn:mavedb:00001201-a). Annotated scoresets are also provided in Table S1.

## Acknowledgments

This work was supported by the Northwestern University – Flow Cytometry Core Facility supported by Cancer Center Support Grant (NCI CA060553). Flow Cytometry Cell Sorting was performed on a BD FACSAria SORP system and BD FACSymphony S6 SORP system, purchased through the support of NIH 1S10OD011996-01 and 1S10OD026814-01. We thank David Liu and the Liu lab for sharing their prime editing reagents. The authors would like to acknowledge Addgene for its invaluable service which facilitated this study. We further thank the Plasmidsaurus sequencing team for making quality control and pilot experiments more feasible and accessible than in the past.

## Contributions

Conceptualisation: GLC, JDC, RJ, DF, LMS. Investigation: CGB, JDC, ZRW, NDC, MKW, SP, AVM, AJV, SD. Visualisation: DLH, JDC. Formal analysis: DLH, NDC, JDC. Data curation: PG, AM, JDW, MER, TC, RS, DZ, PR, MT. Project administration: GLC, RJ, DF, LMS, JDC. Writing—original draft: JDC. Writing—editing: JDC, PG, AM, JDW, MER, TC, NDC, RJ, DLH, LMS, CGB.

