## Supplementary Methods and Figures for "An integrated, scaled approach to resolve TSC2 variants of uncertain significance"

*Cell lines and plasmids (cliPE)*

HEK 293T (ATCC® CRL-3216™) cells were maintained in standard growth medium (Gibco DMEM #11995-073) supplemented with 10% FBS (R&D Systems; 50-152-7067) and PenStrep (Gibco #15140122). HAP1s (Horizon #C669) were maintained in standard growth medium (Gibco IMDM #12440061) supplemented with 10% FBS and PenStrep. HAP1 cells were used at low passage (<10) for all experiments to minimize the percentage of cells with diploid genomes.

pCMV-PEmax-P2A-GFP was a gift from David Liu (Addgene plasmid #180020; http://n2t.net/addgene:180020; RRID:Addgene180020).^21^ pEF1a-hMLH1dn was a gift from David Liu (Addgene plasmid #174824; http://n2t.net/addgene:174824; RRID:Addgene174824).^22^ The pU6-tevopreq1-GG-acceptor was a gift from David Liu (Addgene plasmid #174038; http://n2t.net/addgene:174038; RRID:Addgene174038).^22^ BPK1520 was a gift from Keith Joung (Addgene plasmid #65777; http://n2t.net/addgene:65777; RRID:Addgene65777).^23^ Integrity of all plasmids was confirmed by Sanger or long-read sequencing.

*Cloning individual prime editing constructs (cliPE)*

We screened for active enhanced prime editing guide RNAs (epegRNAs) following Liu lab recommendations.^24^ For the epegRNA screen, a single epegRNA for each exon was cloned into the pU6-tevopreq1-GG-acceptor backbone using standard Golden Gate assembly (Table S6).^25^ For the MAVE, a nicking gRNA-encoding plasmid for each target exon was cloned using an adjusted version of the Zhang lab target sequence cloning protocol.^26^ Briefly, oligonucleotides encoding the nicking gRNA were synthesized (IDT; Table S5), annealed and phosphorylated, and ligated into linearized expression vector BPK1520.

*Cloning epegRNA libraries (cliPE)*

Pools of single-stranded DNA oligos encoding epegRNAs were designed and ordered (IDT oPools; Table S3). The oligo pool was made double-stranded and BsaI restriction enzyme recognition sites were appended via PCR (10 cycles). The PCR-amplified oligo pool was then digested with BsaI-HFv2 (NEB #R3733S), according to the manufacturer’s instructions. The destination vector (pU6-tevopreq1-GG-acceptor) was also linearized by restriction digest with BsaI-HFv2 and purified by gel or column purification. The digested oligos and tevopreq vector were ligated using T4 DNA ligase (NEB #M0202S) according to the manufacturer’s instructions. The pool of assembled plasmids was then transformed into OneShot TOP10 chemically competent *E. coli* (Thermo Fisher #C404003). The transformation was plated onto a 10-cm LB agar plate containing ampicillin (100 µg/mL). The bacteria were cultured overnight before the plates of colonies were scraped. The ZymoPURE II plasmid midiprep kit (Fisher #NC0835048) was used to extract plasmid DNA from the pooled colonies. Once library concentrations were determined, plasmid libraries were analyzed via long-read sequencing (Plasmidsaurus) and custom targeted amplicon sequencing for quality control.

*Prime editing in HEK293Ts (cliPE)*

The co-transfection of the epegRNA-containing pU6-tevopreq1-GG-acceptor, pCMV-PEmax-P2A-GFP, and pEF1a-hMLH1dn plasmids into HEK293T cells was carried out using the TurboFectin 8.0 transfection reagent (OriGene #TF81005). 100,000 cells were seeded 24 hours before transfection in 24-well plates. Plasmids were co-transfected in the following amounts: 75 ng (pU6-tevopreq1-GG-acceptor), 263 ng (pCMV-PEmax-P2A-GFP), and 132 ng (pEF1a-hMLH1dn). As the PEmax plasmid also encodes GFP, successfully transfected cells were selected using fluorescence-activated cell sorting (FACS). GFP+ cells were re-plated under standard culture conditions for at least 48 hours. Cell pellets were either used immediately for genomic DNA (gDNA) extraction or frozen at –20°C prior to gDNA extraction. gDNA from prime edited cells was screened for editing using PCR amplifying the target exon followed by Sanger sequencing (Table S4).

*Prime editing in haploid HAP1s (cliPE)*

The co-transfection of the epegRNA-containing pU6-tevopreq1-GG-acceptor library, pCMV-PEmax-P2A-GFP, pEF1a-hMLH1dn, and nicking gRNA-containing BPK1520 plasmids into HAP1 cells was carried out using the Neon electroporation system (Thermo Fisher #MPK5000). For co-transfection, 1 million HAP1 cells were electroporated with the following amounts of plasmid DNA (total of 3000 ng): 451 ng (pU6-tevopreq1-GG-acceptor), 1579 ng (pCMV-PEmax-P2A-GFP), 789 ng (pEF1a-hMLH1dn), and 180 ng (BPK1520). Successfully transfected cells were selected using FACS sorting for GFP+ cells. GFP+ cells (typically 15-30%) were re-plated under standard culture conditions until at least 1 million cells were obtained.

*pS6 FACS (cliPE)*

HAP1 cells were washed twice with DPBS and serum-starved overnight by incubation in low-sera media (IMDM + 0.2% FBS). Serum starvation has been previously utilized in low-throughput experiments to establish that loss of *TSC2* function results in constitutive mTORC1 activity.^27,28^ Starved HAP1 cells were washed twice with DPBS supplemented with 1% v/v Phosphatase Inhibitor Cocktail 3 (Sigma #P0044) before trypsinization with trypLE (trypLE; 5 min; Gibco #25300062) supplemented with 1% v/v Phosphatase Inhibitor Cocktail 3. Cells were then centrifuged and resuspended in BD Fixation/Permeabilization solution (BD Biosciences #554714). After incubating in the fixation solution on ice for 20 minutes, the HAP1 cells were washed twice with BD permeabilization/wash buffer. The CST Alexa594 conjugated rabbit anti-phospho-S6 antibody (#5018; D68F8) was then used to immunolabel pS6. Cells were incubated with the antibody (1:50 dilution) for 30 minutes on ice in the dark. This antibody had been previously validated for the functional characterization of variants in mTORopathy gene *SZT2*.^29^ Immunolabeled cells were then washed twice more with BD permeabilization/wash buffer before they were flow sorted by pS6 level on the BD FACSMelody 3-laser cell sorter. A minimum of 200,000 cells in the upper quartile for pS6 were collected. Unsorted cells were also reserved for downstream processing.

*Amplicon sequencing and data analysis (cliPE)*

The gDNA of the fixed, FACS-sorted HAP1 cells was extracted using the PureLink™ Genomic DNA Mini Kit (Invitrogen #K182002). Manufacturer’s instructions were followed with the exception of an added incubation (40-60 minutes, 90°C) to reverse formaldehyde-induced crosslinks. The target regions of *TSC2* were then amplified via PCR (Bio-Rad iProof system), followed by an additional barcoding PCR reaction. The locus was then sequenced via short-read amplicon sequencing (Illumina MiniSeq System; 100,000 minimum read-depth). Reads were aligned to the human genome (hg38) and variant allele frequencies were calculated using the jellyfish k-mer counting package.^30^ Variants with allele frequency below 0.1% in unsorted cells were filtered out. Variant allele frequencies were then used to calculate a TSC2 activity score. All cliPE scores are reported in Table S1.

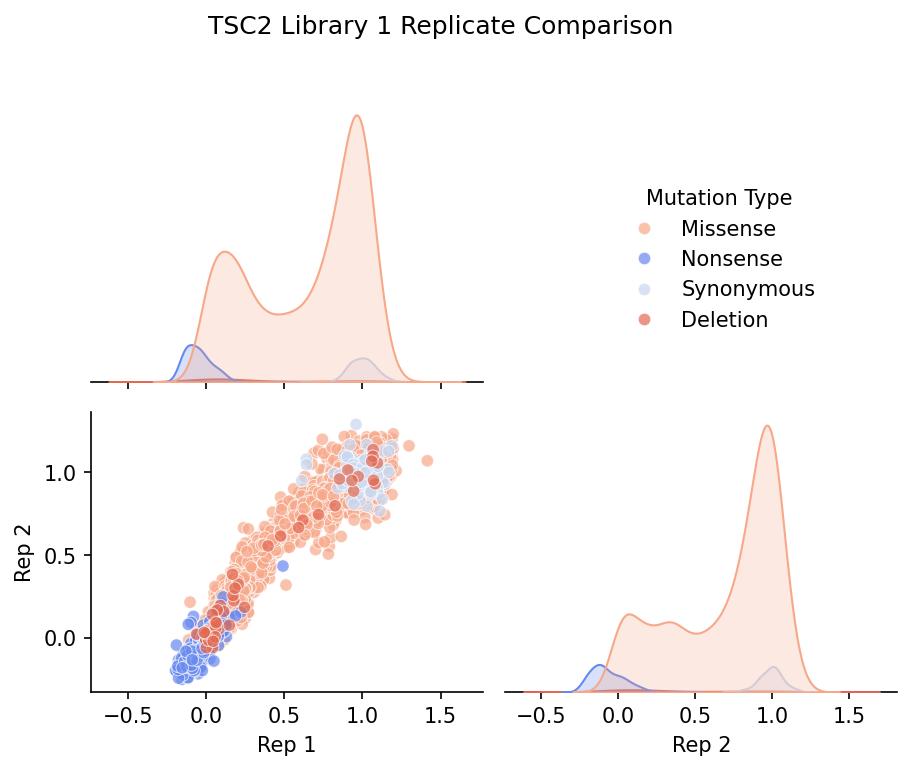

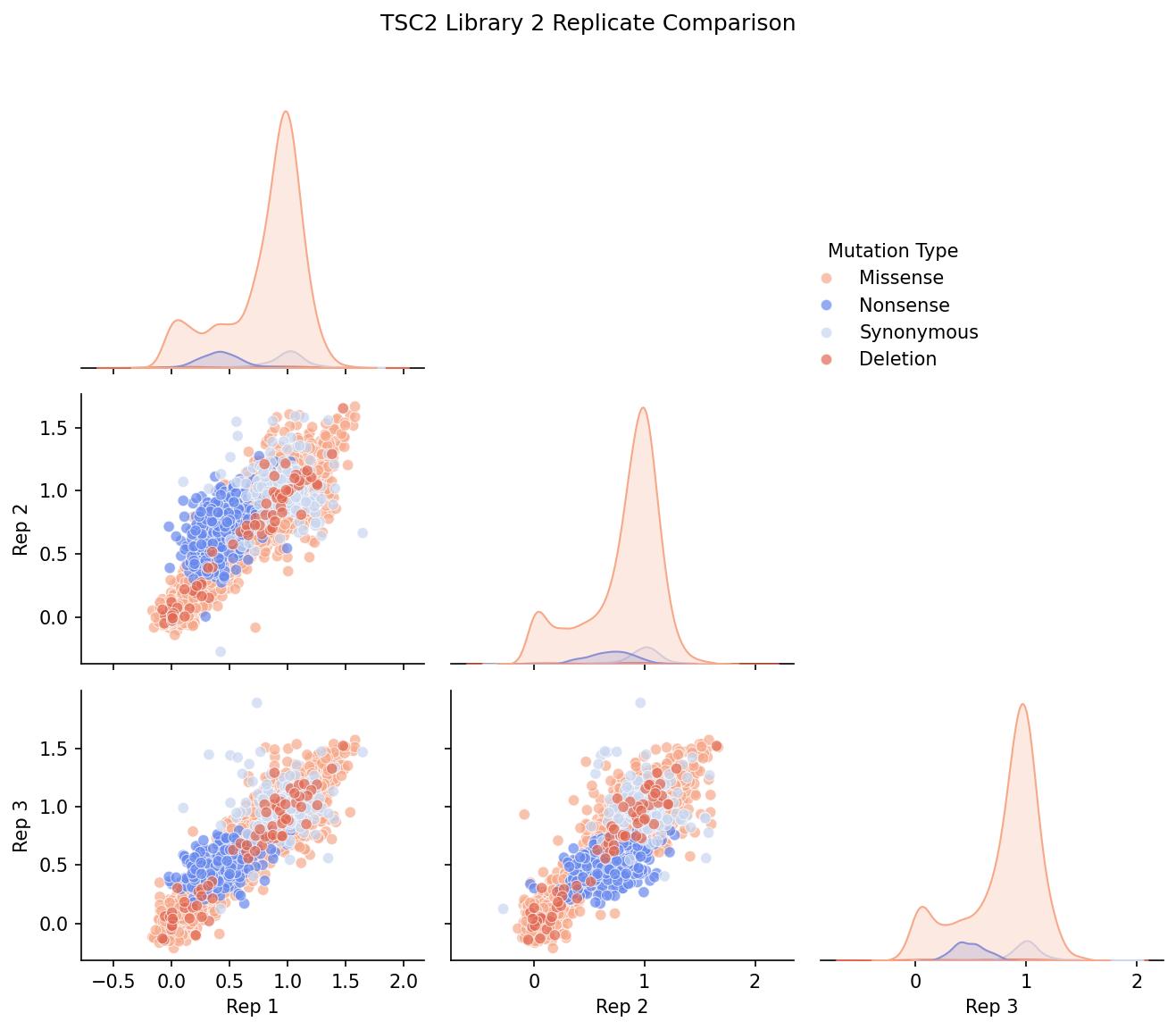

### Figure S1: Replicate correlation plots for VAMP-seq.

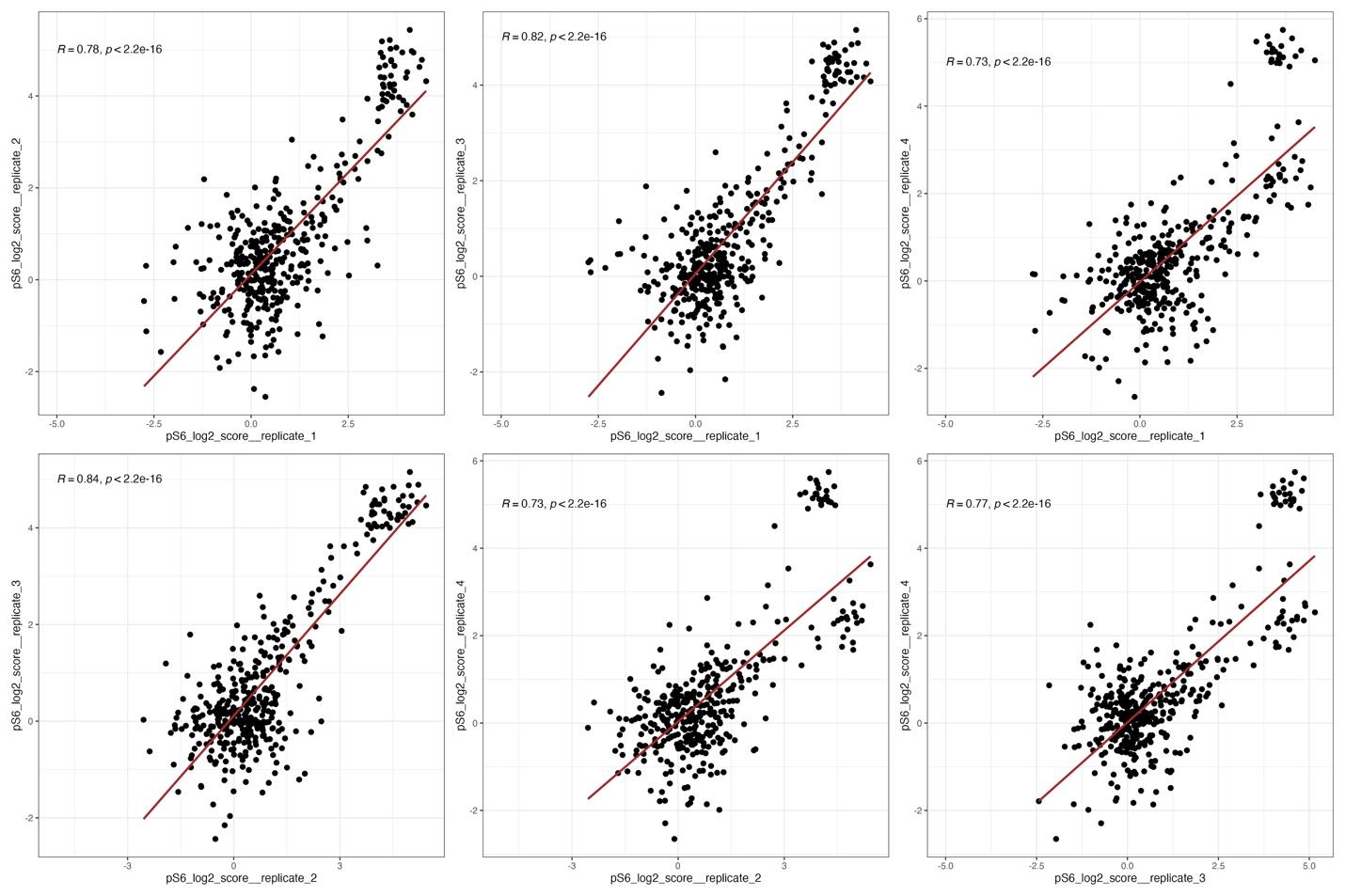

### Figure S2: Replicate correlation plots for immune SGE.

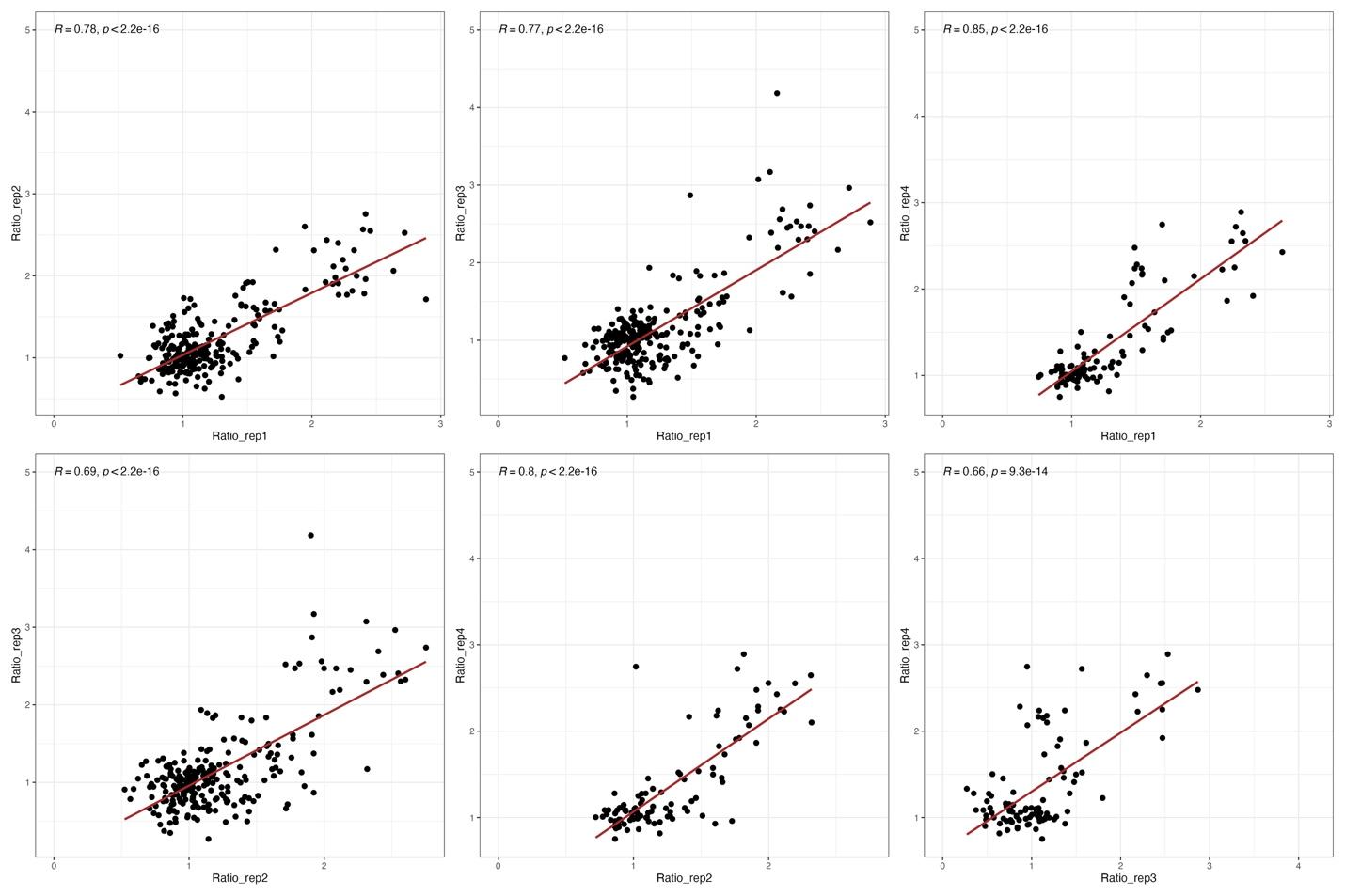

### Figure S3: Replicate correlation plots for cliPE.

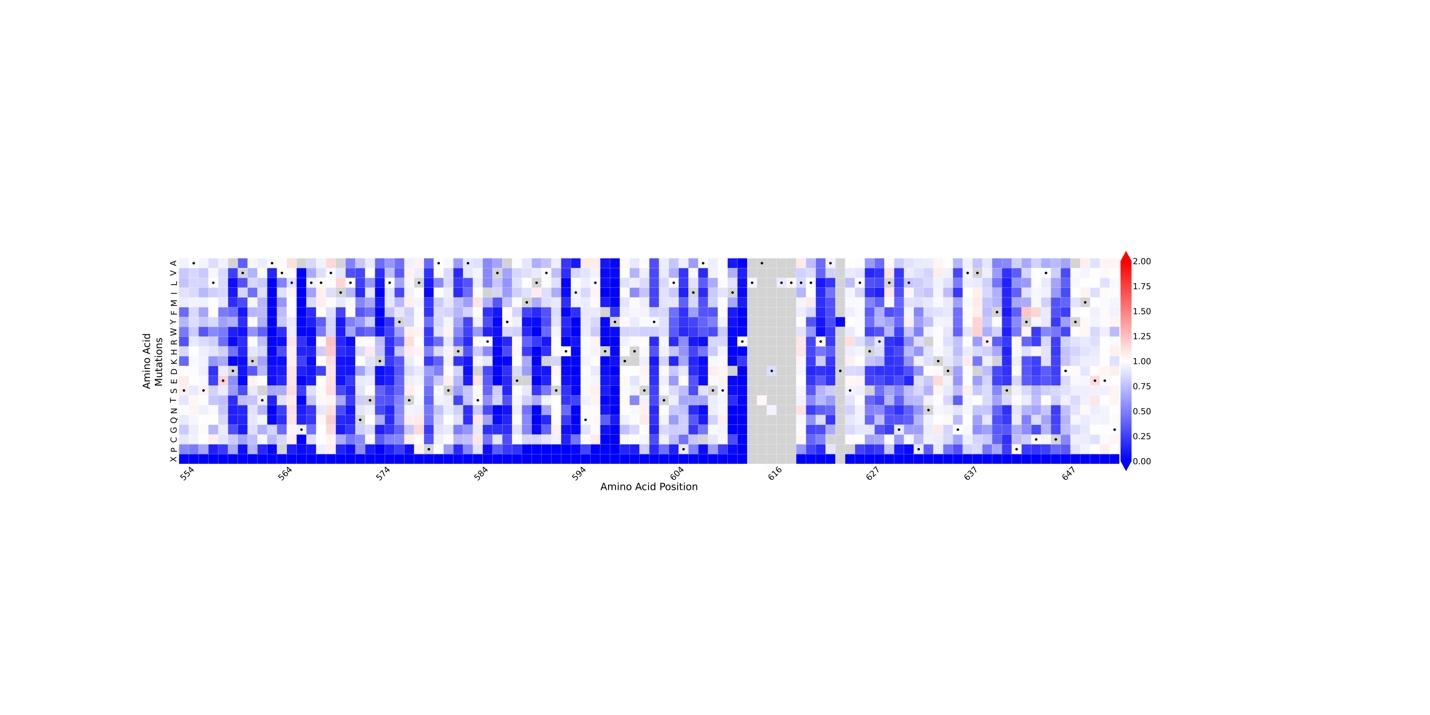

### Figure S4: Heatmap representation of TSC2 abundance VAMP-seq dataset for amino acid residues 554 to 654.

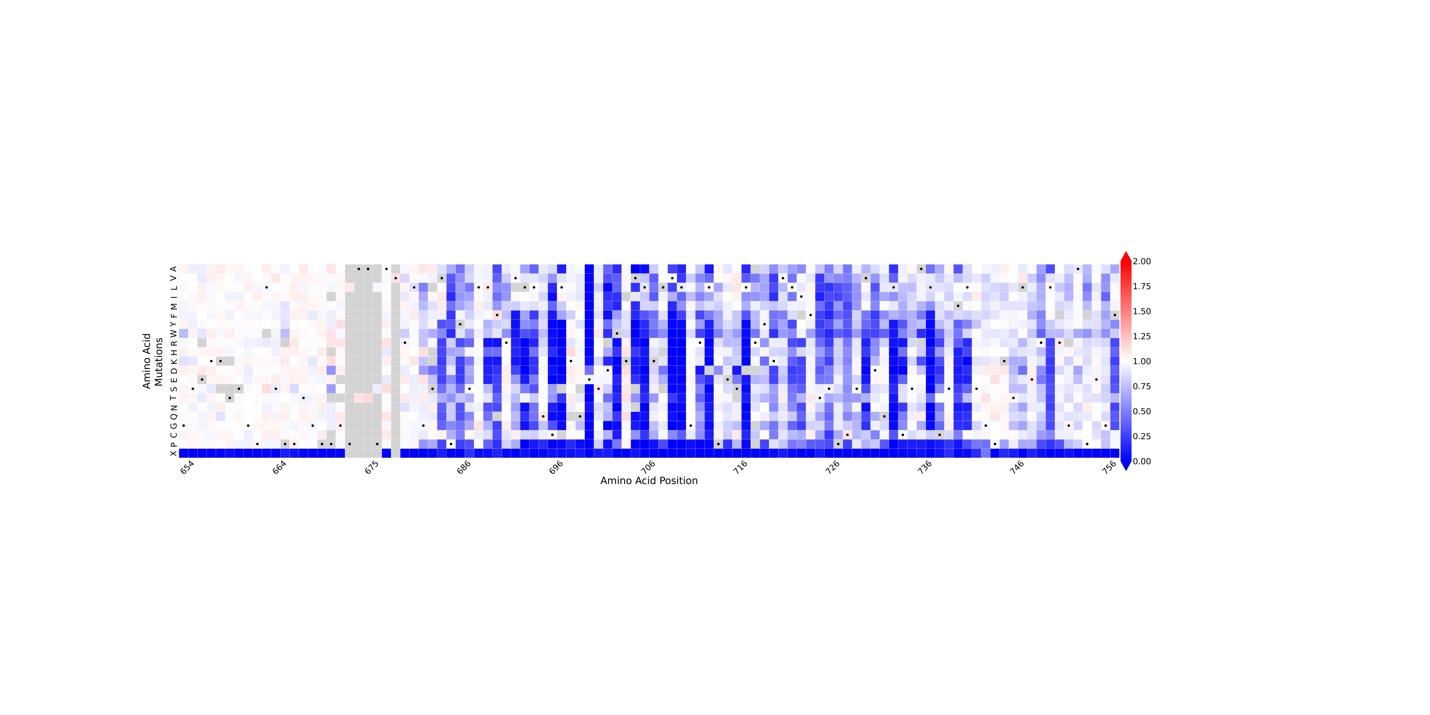

### Figure S5: Heatmap representation of TSC2 abundance VAMP-seq dataset for amino acid residues 654 to 757.

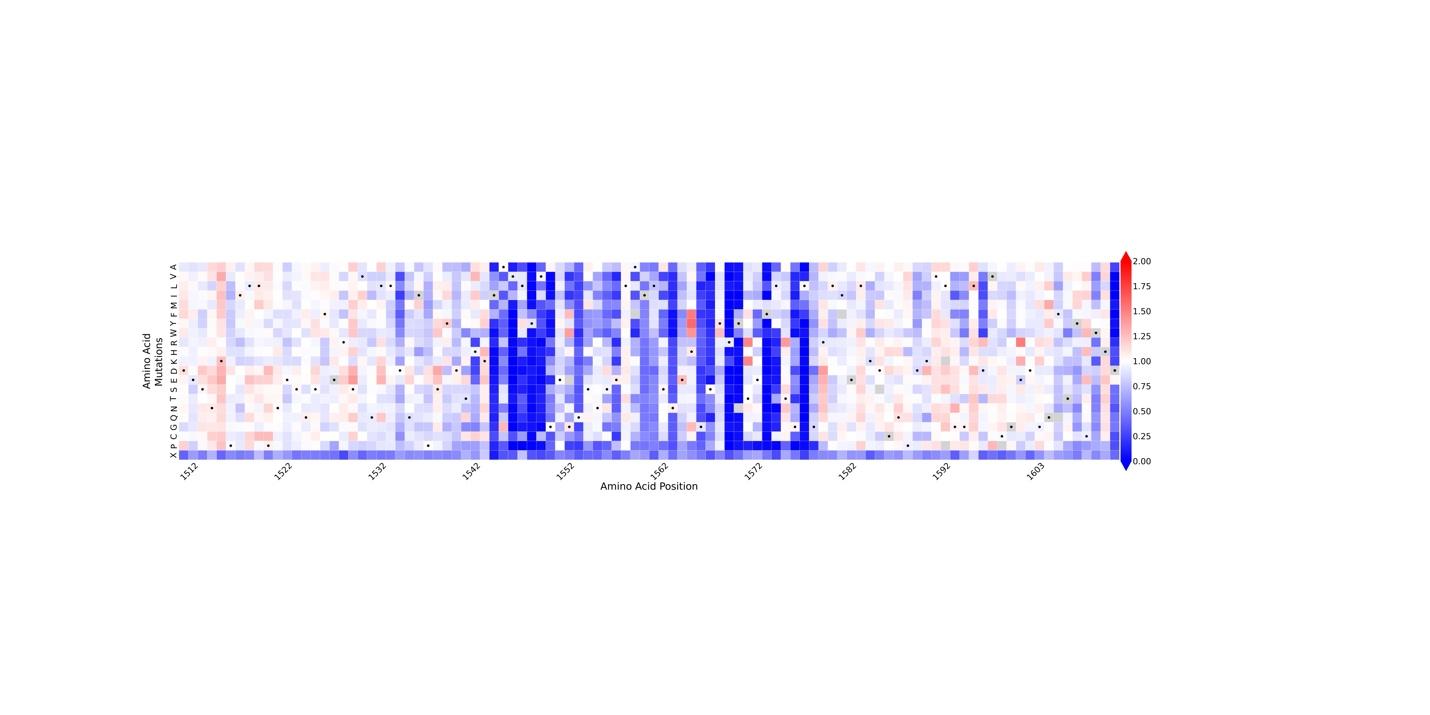

### Figure S6: Heatmap representation of TSC2 abundance VAMP-seq dataset for amino acid residues 1512 to 1612.

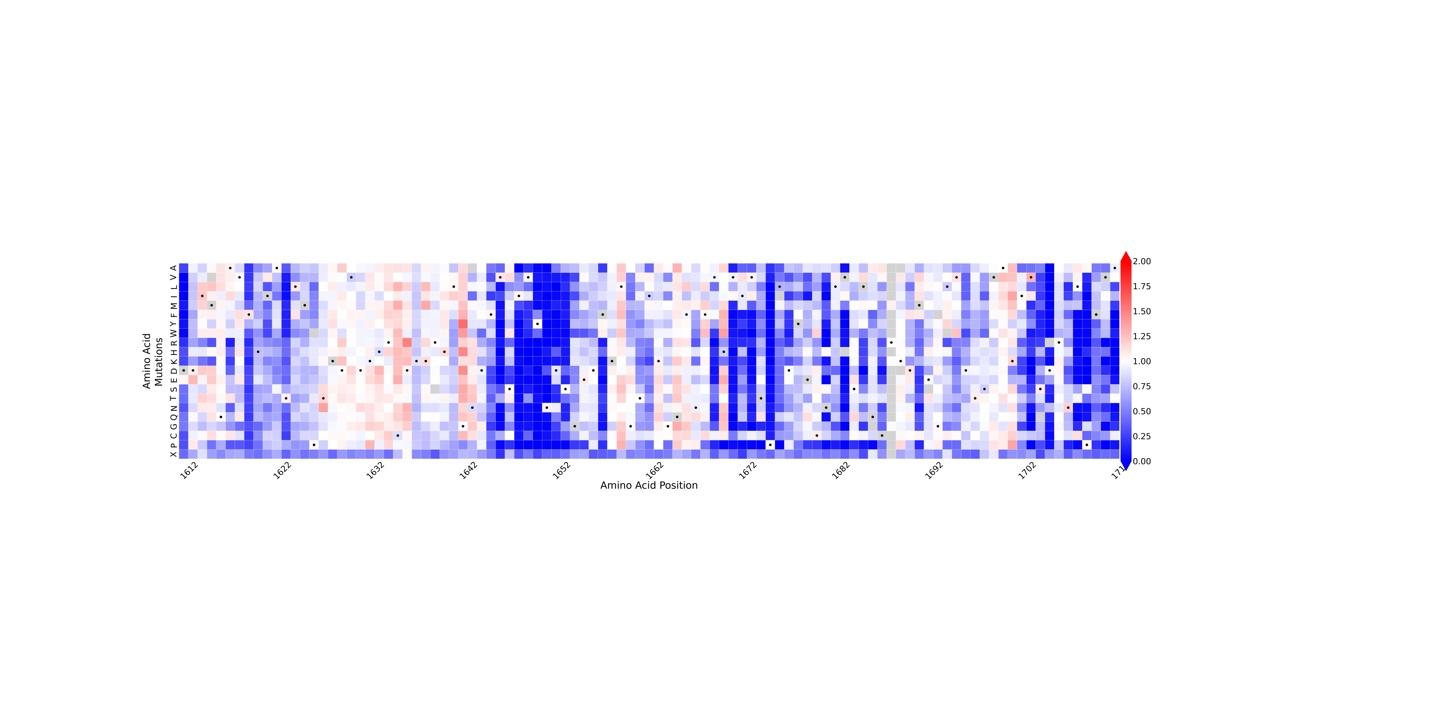

### Figure S7: Heatmap representation of TSC2 abundance VAMP-seq dataset for amino acid residues 1612 to 1712.

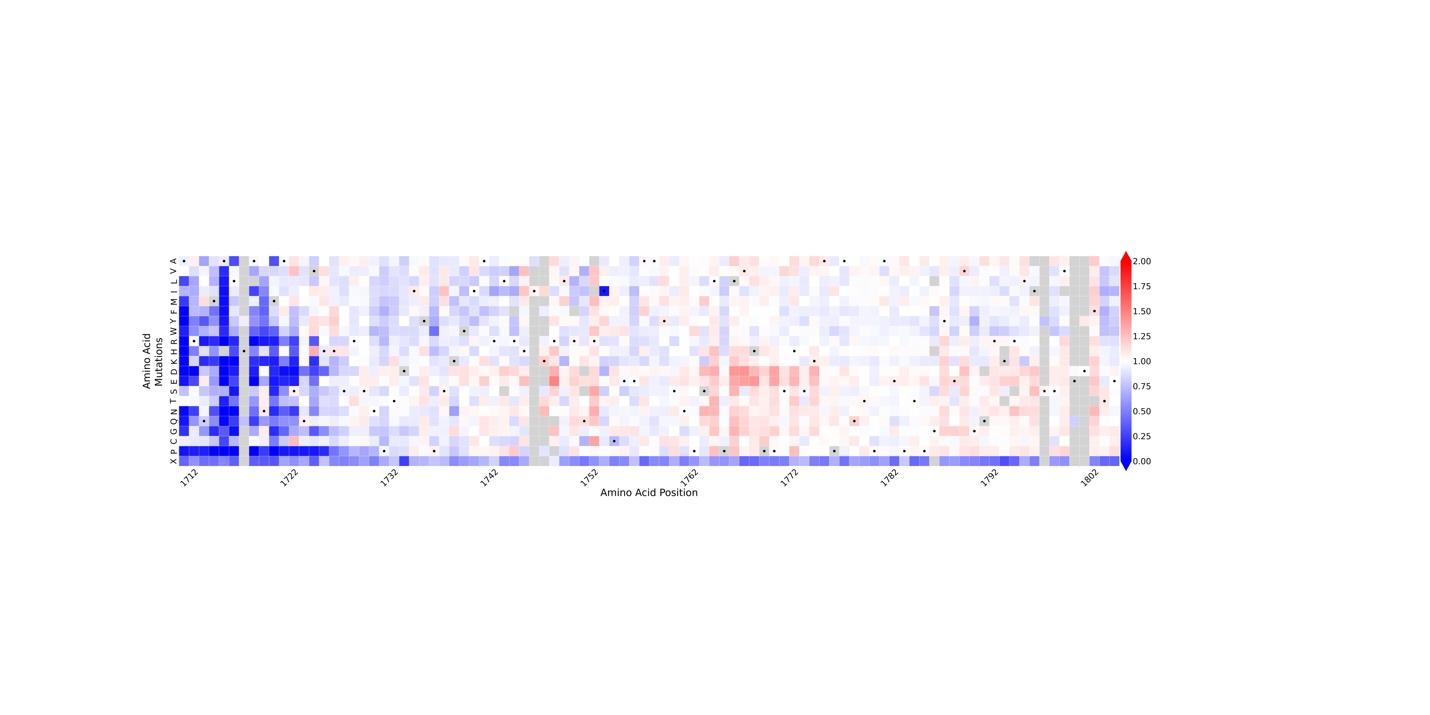

### Figure S8: Heatmap representation of TSC2 abundance VAMP-seq dataset for amino acid residues 1712 to 1805.

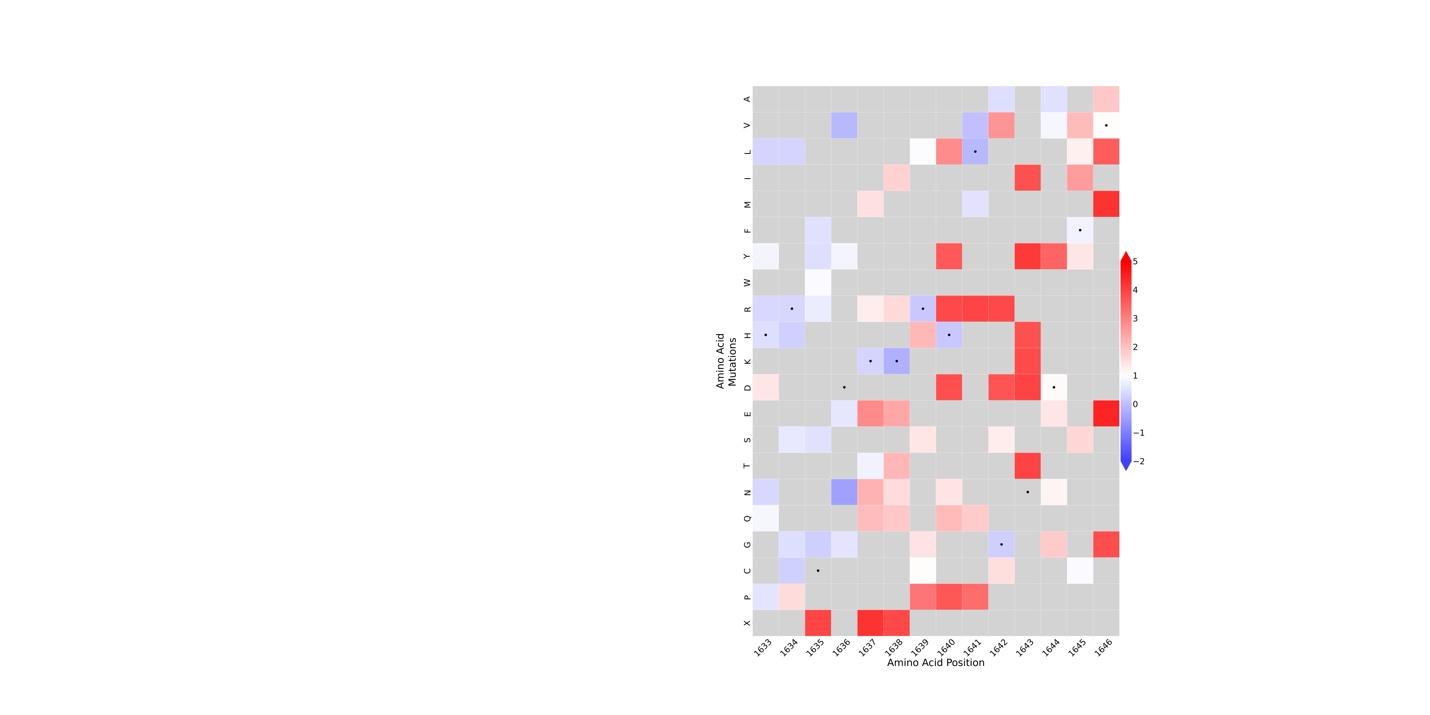

### Figure S9: Heatmap representation of TSC2 function immune SGE dataset for amino acid residues 1633 to 1646.

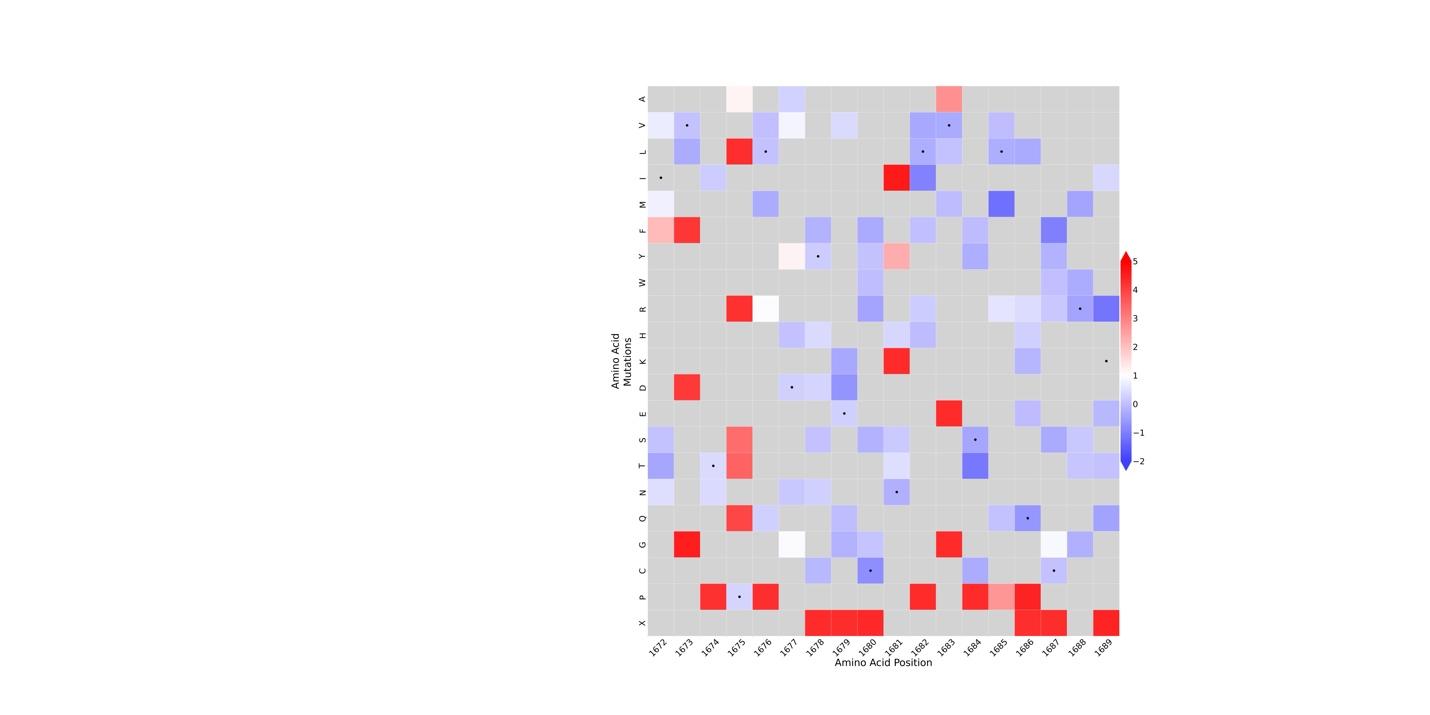

### Figure S10: Heatmap representation of TSC2 function immune SGE dataset for amino acid residues 1672 to 1689.

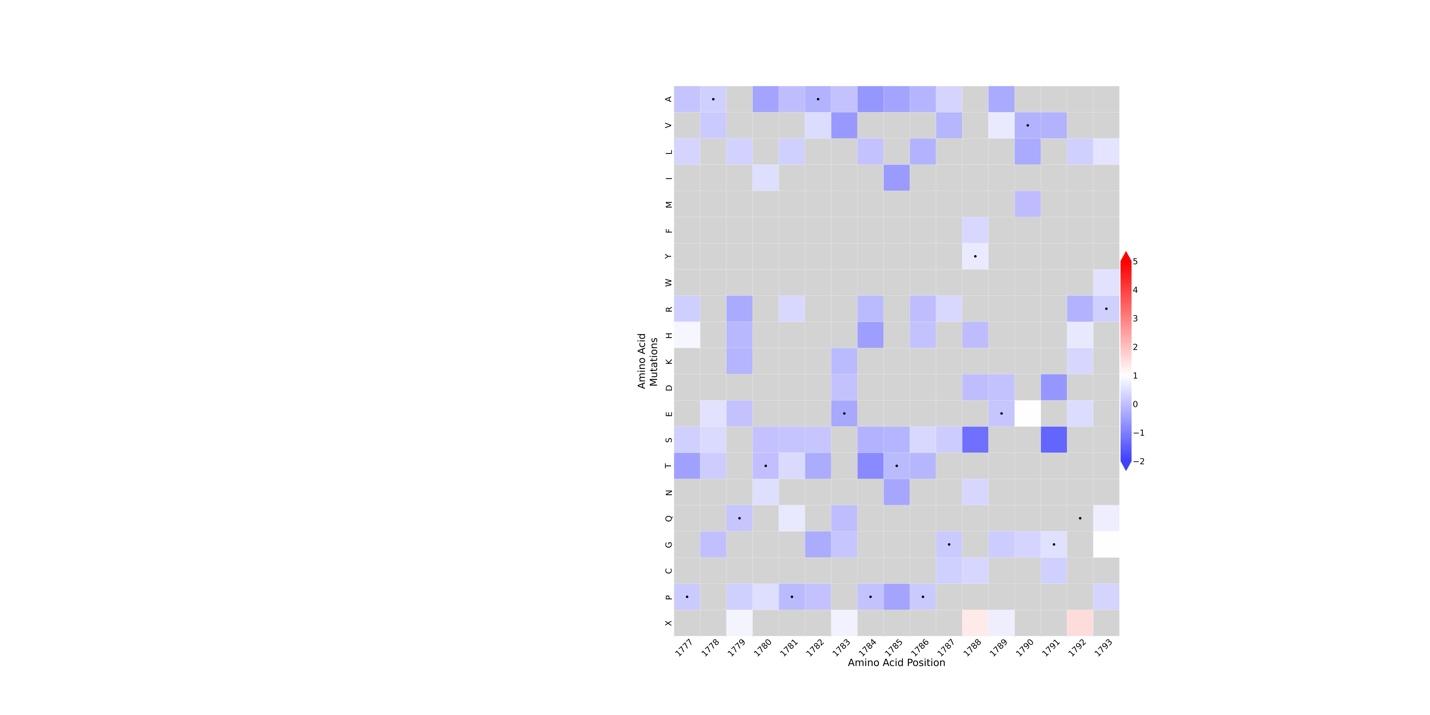

### Figure S11: Heatmap representation of TSC2 function immune SGE dataset for amino acid residues 1777 to 1793.

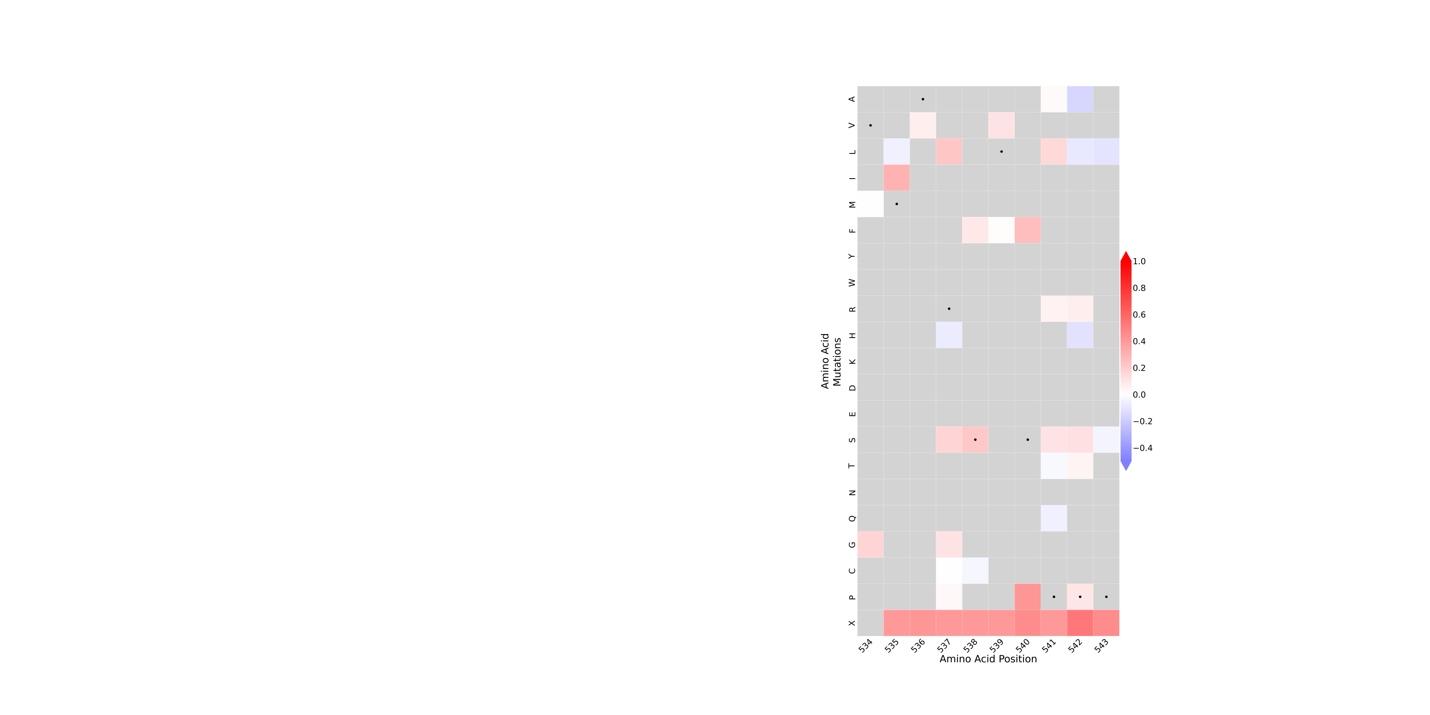

### Figure S12: Heatmap representation of TSC2 function cliPE dataset for amino acid residues 534 to 543.

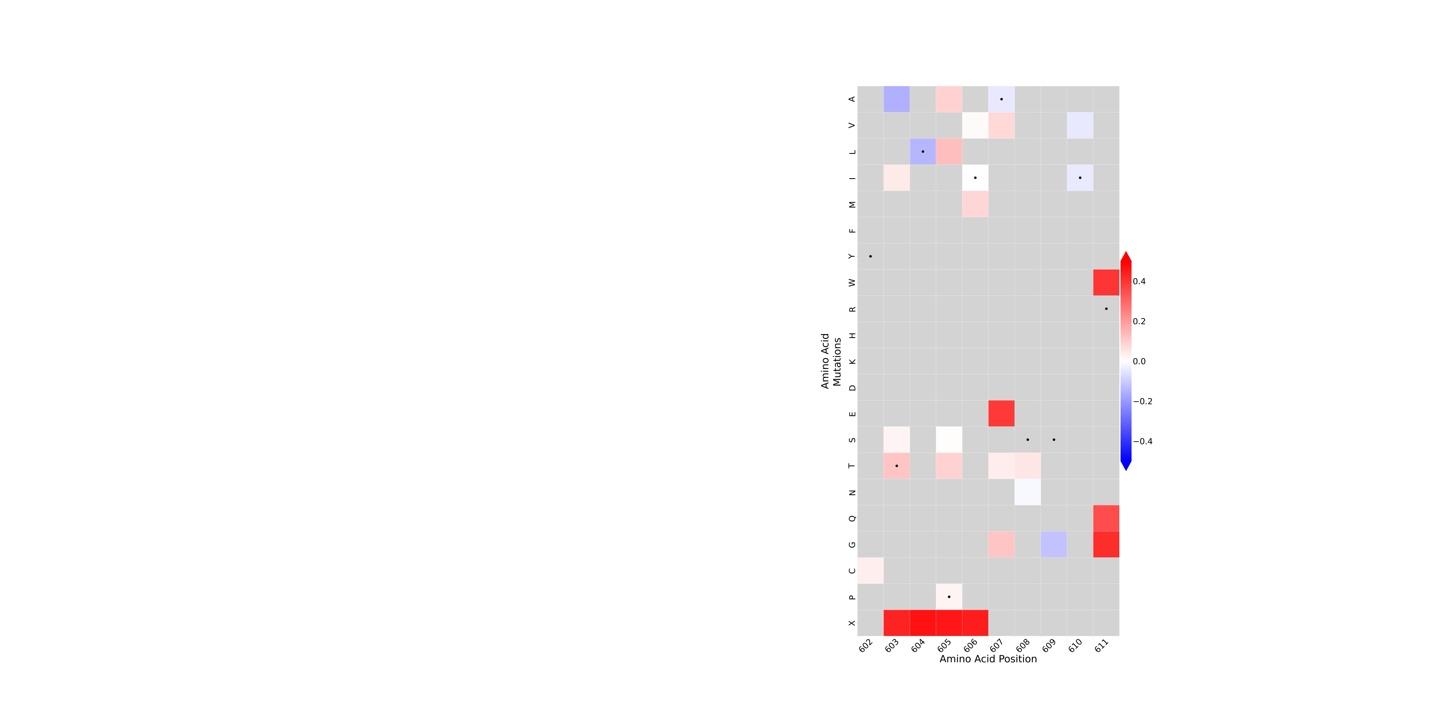

### Figure S13: Heatmap representation of TSC2 function cliPE dataset for amino acid residues 602 to 611.

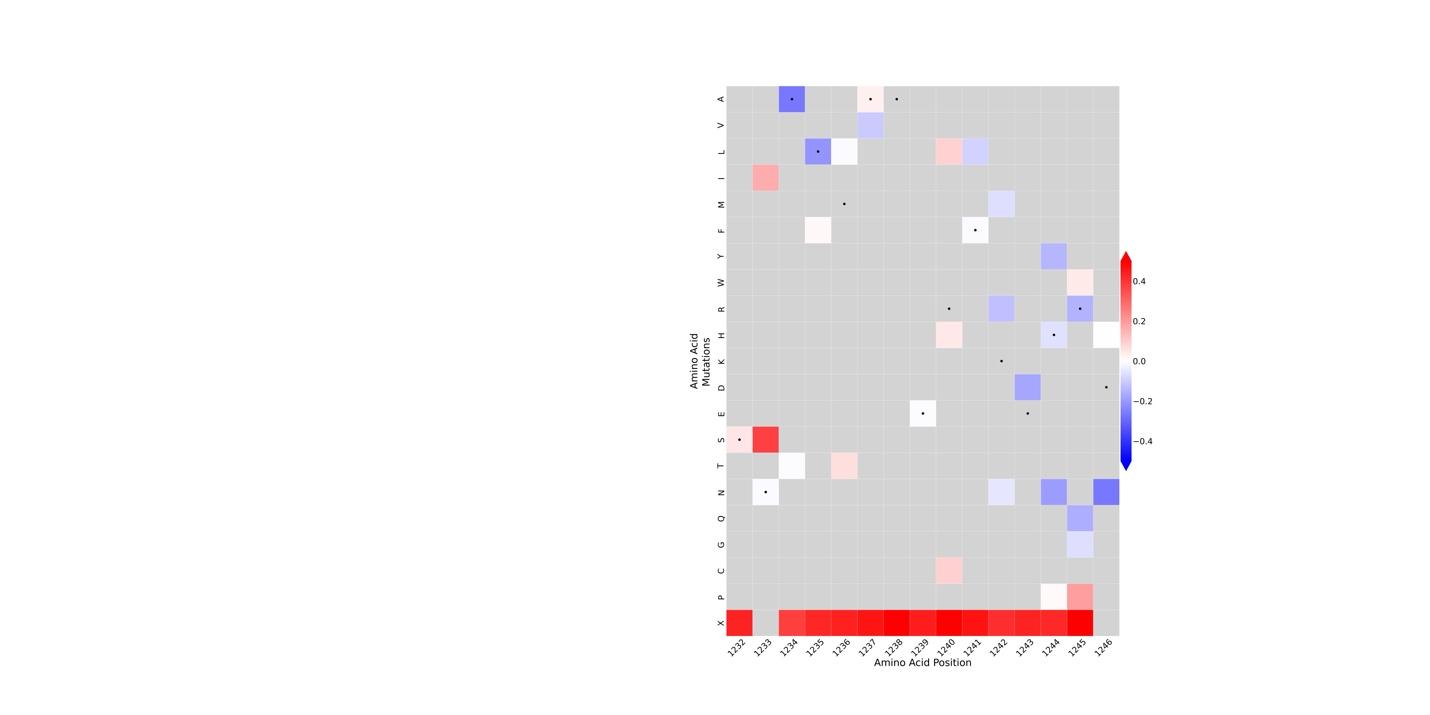

### Figure S14: Heatmap representation of TSC2 function cliPE dataset for amino acid residues 1232 to 1246.

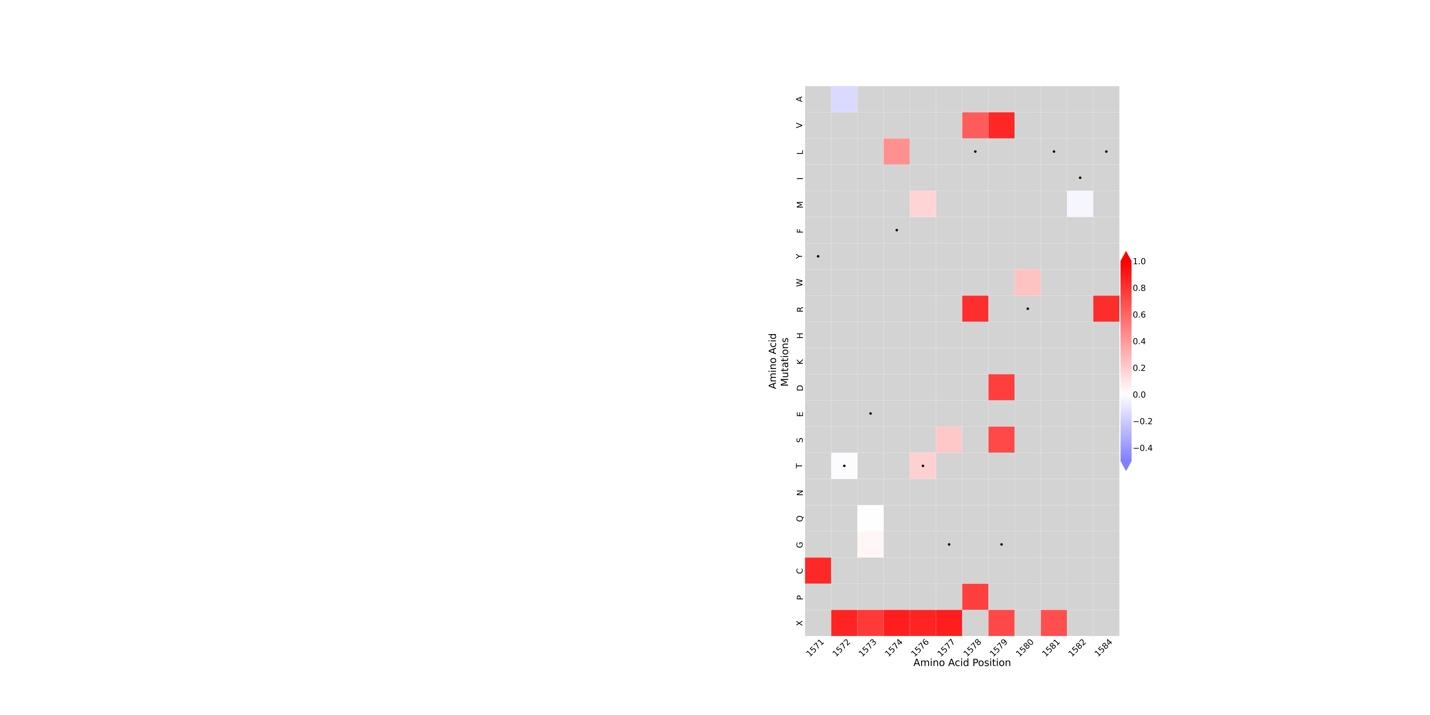

### Figure S15: Heatmap representation of TSC2 function cliPE dataset for amino acid residues 1571 to 1584.

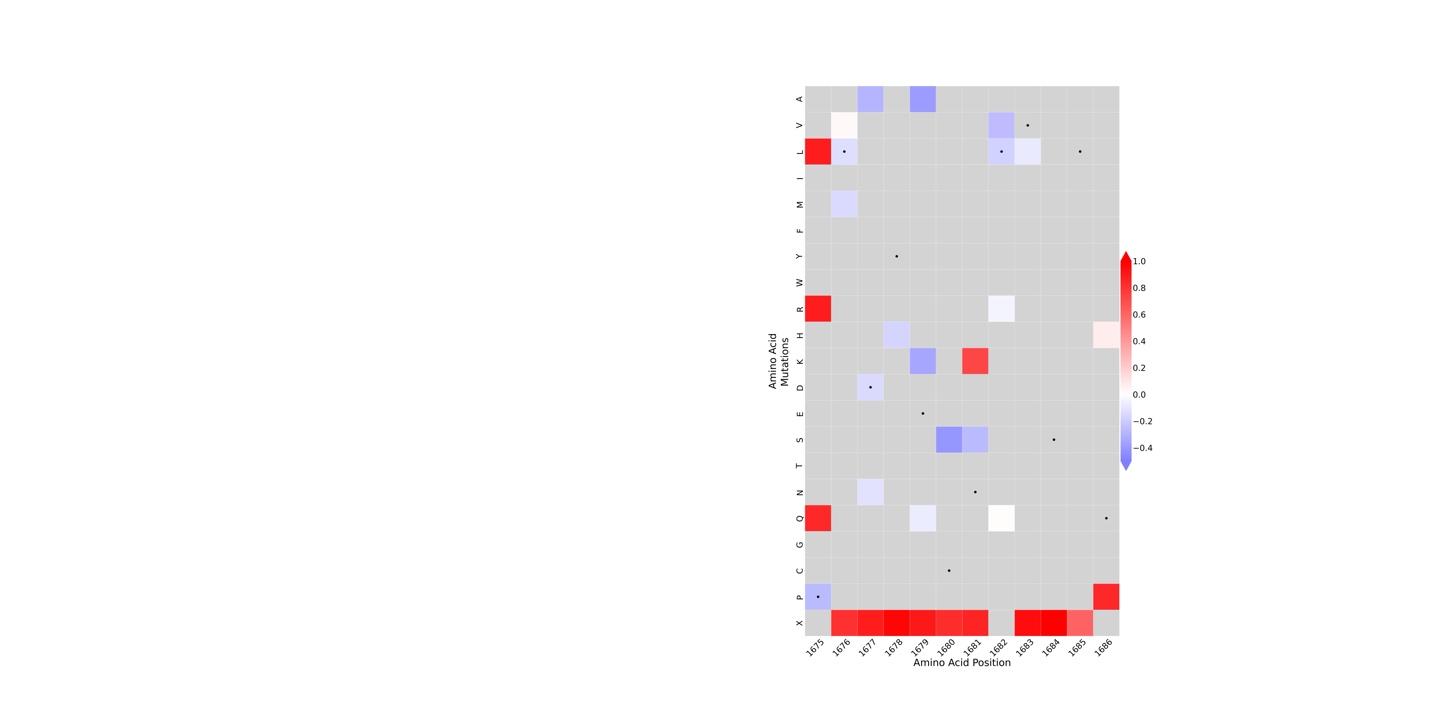

### Figure S16: Heatmap representation of TSC2 function cliPE dataset for amino acid residues 1675 to 1686.

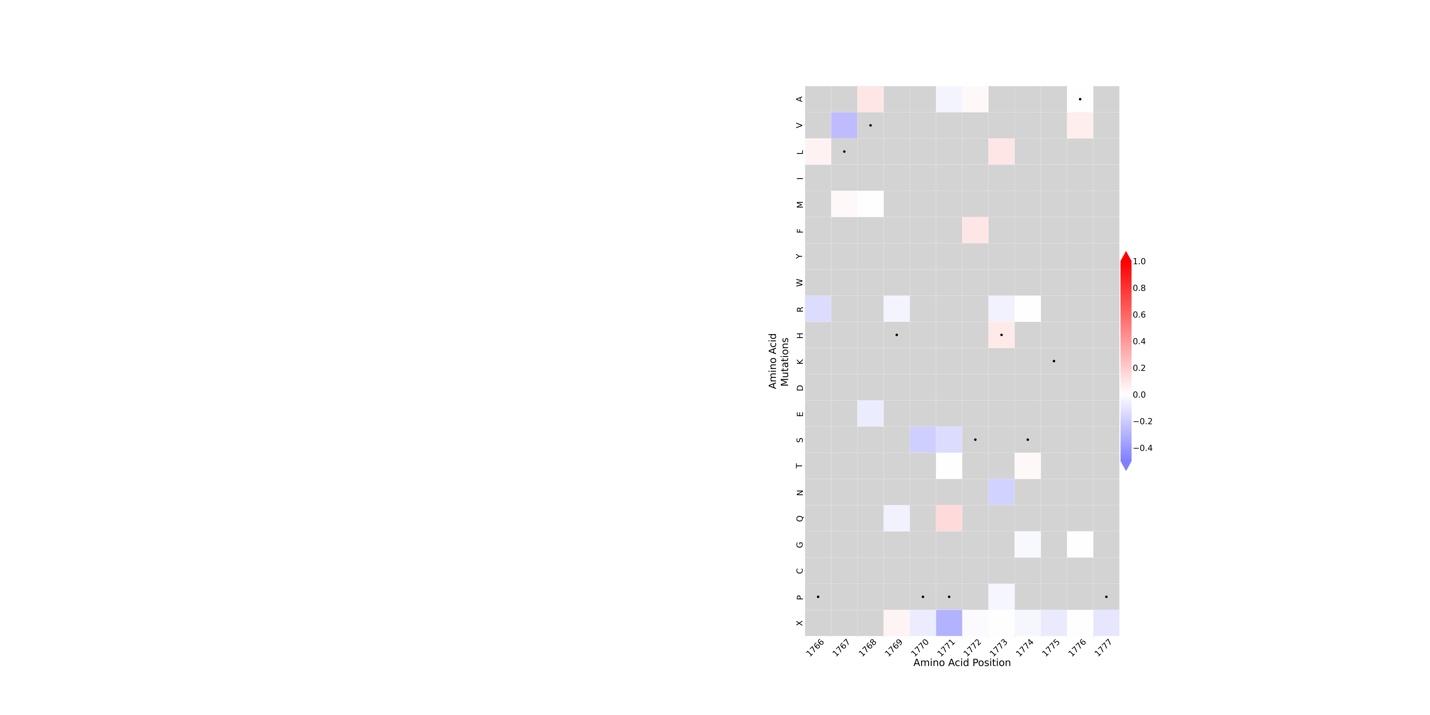

### Figure S17: Heatmap representation of TSC2 function cliPE dataset for amino acid residues 1766 to 1777.

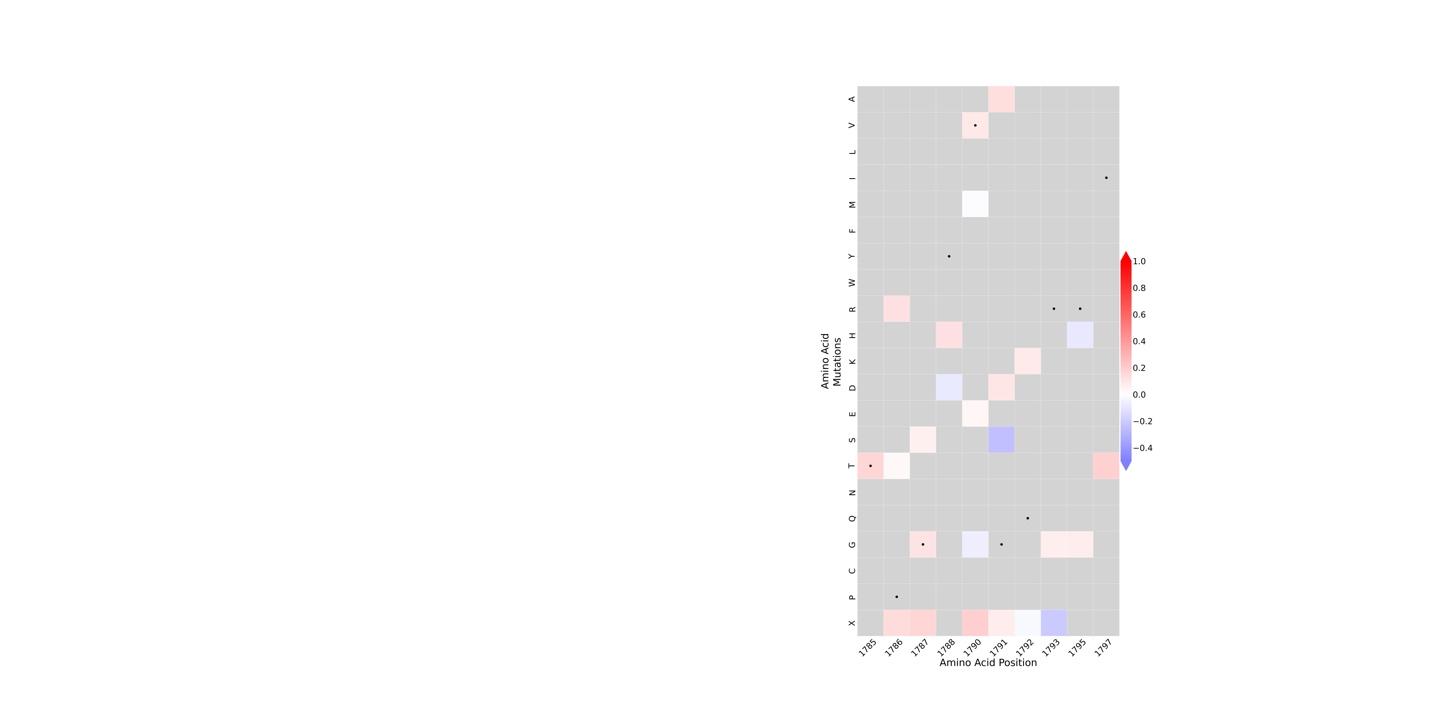

### Figure S18: Heatmap representation of TSC2 function cliPE dataset for amino acid residues 1785 to 1797.

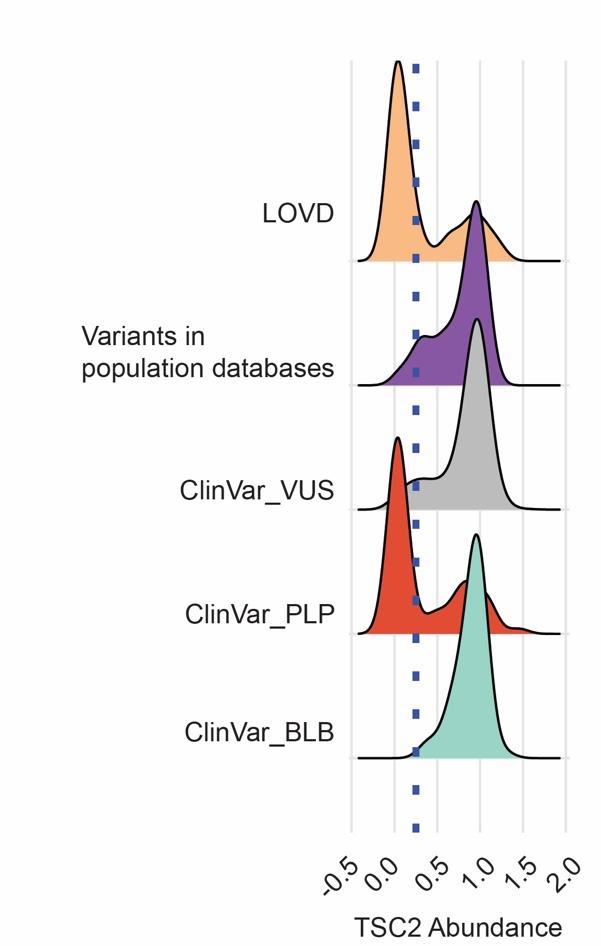

Figure S19: Distribution of TSC2 abundance scores derived from VAMP-seq for distinct subpopulations of variants. ‘LOVD’ = variants described in the Leiden Open Variation Database as pathogenic or likely pathogenic which are distinct from the ClinVar database list of pathogenic or likely pathogenic variants. ‘Variants in population databases’ = missense variants present in either the gnomAD database or the Regeneron million exome database with an allele count >2.

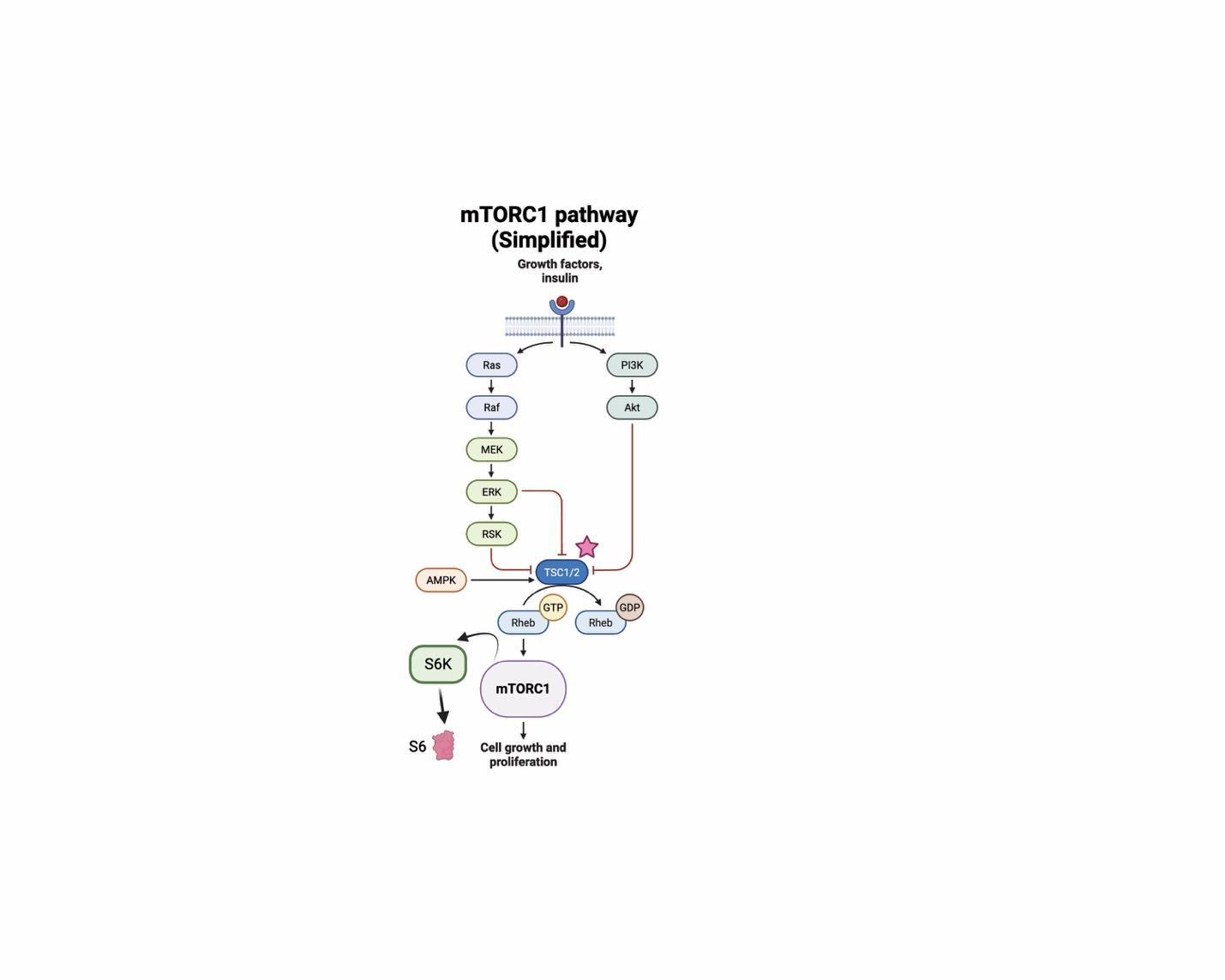

Figure S20: Simplified diagram of the mTORC1 signaling pathway. The TSC1/2 complex acts as a negative regulator of mTORC1 activity. In tuberous sclerosis, the absence of functional TSC1/2 complex results in constitutive activation of mTORC1.

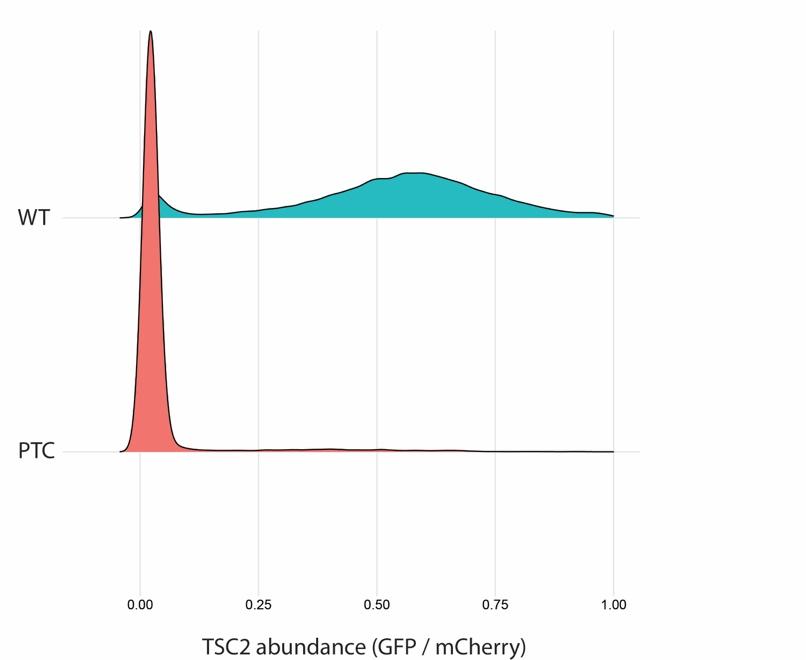

Figure S21: Distributions of TSC2 abundance scores from pseudoclonal cell populations expressing either WT or a premature termination codon (PTC) variant. TSC2-GFP fluorescence was normalized to mCherry fluorescence.

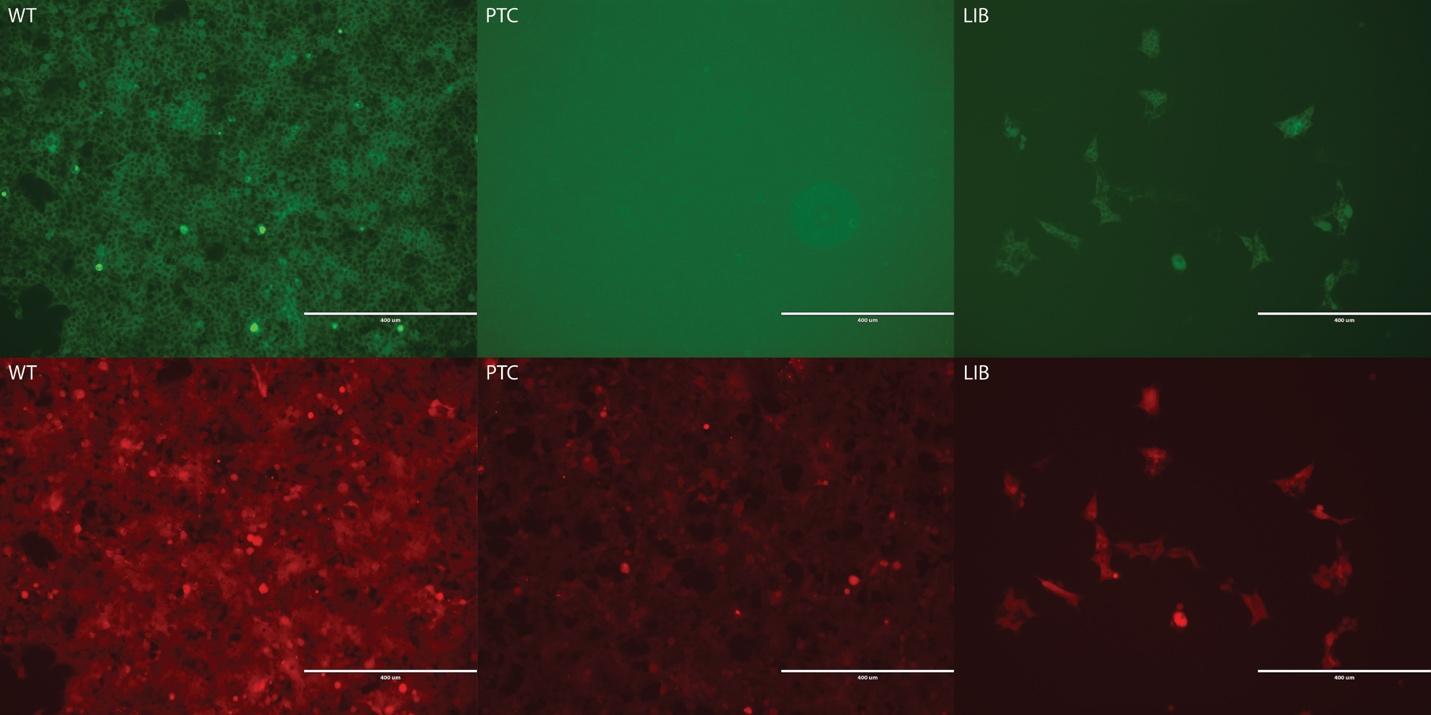

Figure S22: Example images of pseudoclonal HEK landing pad cells after integration of VAMP-seq constructs and selection by rimiducid followed by puromycin. Images captured on benchtop EVOS imaging system with 10X objective. (Top) TSC2-GFP fluorescence. (Bottom) mCherry fluorescence. Abbreviations: “WT” = the WT reference TSC2 cDNA; “PTC” = a representative premature termination codon TSC2 variant; “LIB” = an example of a variant library.

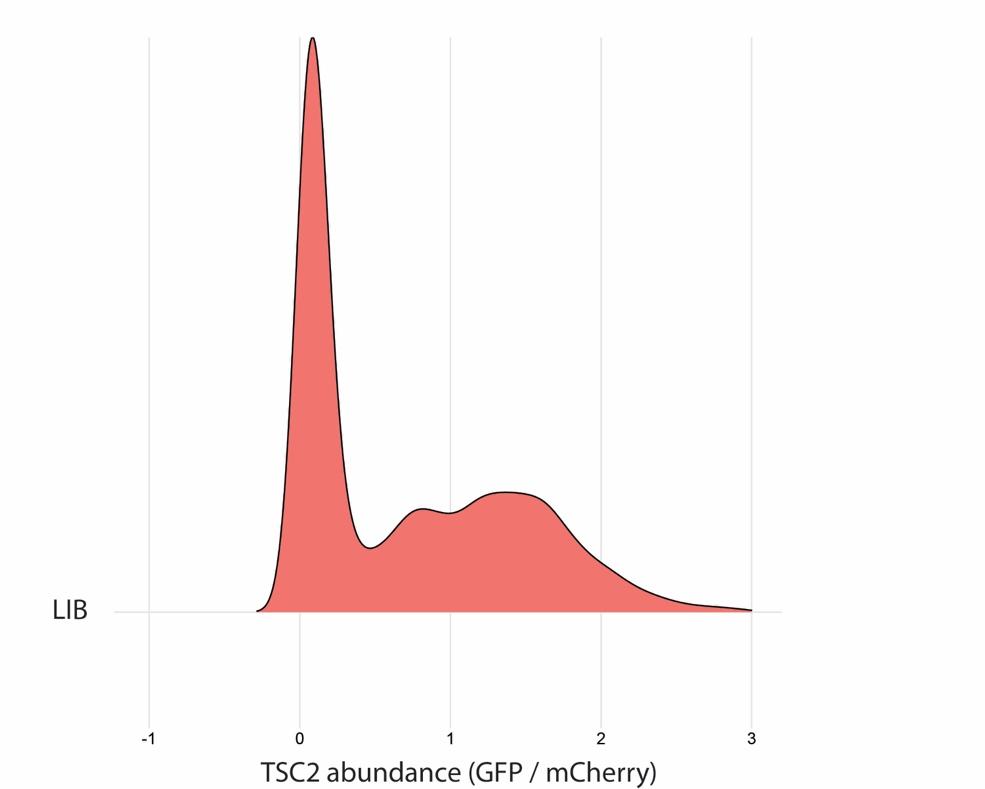

Figure S23: Distributions of TSC2 abundance scores from pseudoclonal cell populations expressing a variant library. TSC2-GFP fluorescence was normalized to mCherry fluorescence.

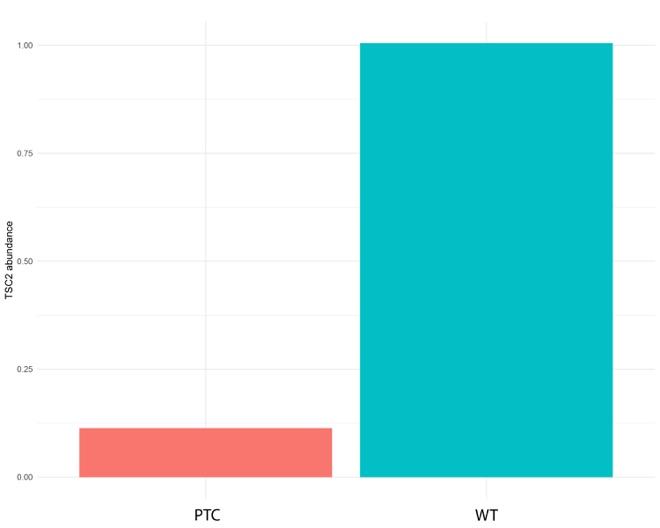

Figure S24: Amplicon sequencing validation of enrichment of WT TSC2-GFP in high GFP bins. HEK landing pad cells expressing a variant library were sorted into 4 bins based on GFP fluorescence. After amplicon sequencing, abundance scores were calculated. WT TSC2-GFP has a TSC2 abundance score of approximately 1, indicating enrichment in high GFP bins. We averaged the abundance score for the three PTC variants with most read-depth coverage. PTC variants had an enrichment score below 0.25, suggesting enrichment in low GFP bins.

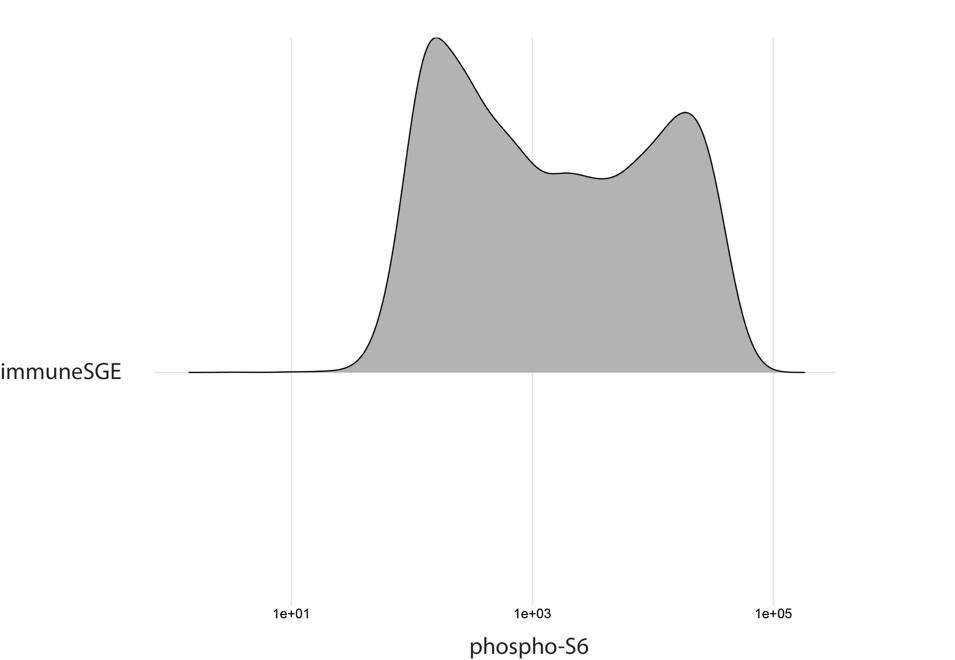

Figure S25: Distribution of phospho-S6 immunolabeling in primary immune CD4+ T cells with saturation genome editing of the TSC2 locus. The top (pS6-HIGH) and bottom (pS6-LOW) quartiles were sorted by FACS.

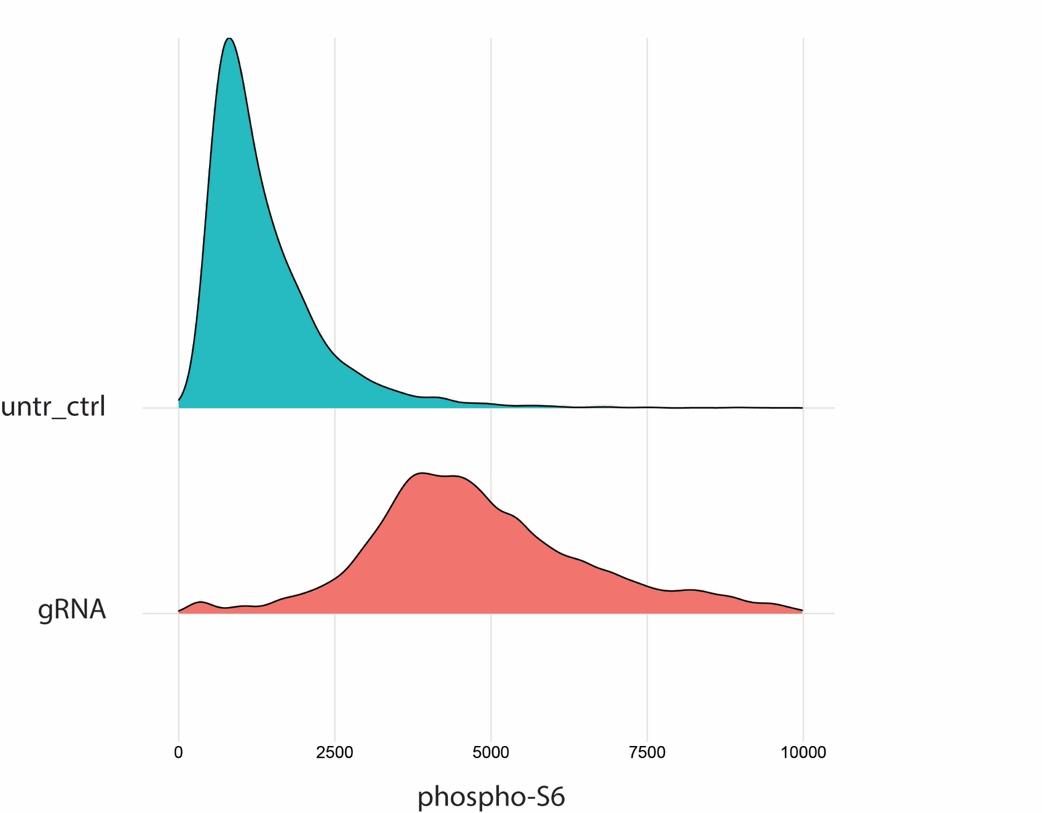

Figure S26: Assay development for cliPE. HAP1 cells were transfected with a gRNA targeting exon 17 of TSC2 cloned into a px458/9 family of vectors to co-express Cas9. We observed elevated levels of pS6 in serum-starved, guide-RNA transfected cells relative to serum-starved untransfected cells. We further validated the generation of NEHJ-induced indels by PCR amplification of exon 17 and sequencing amplicons (Figure S27). Abbreviations: ‘untr_ctrl’ = untransfected control; ‘gRNA’ = gRNA transfected cells.

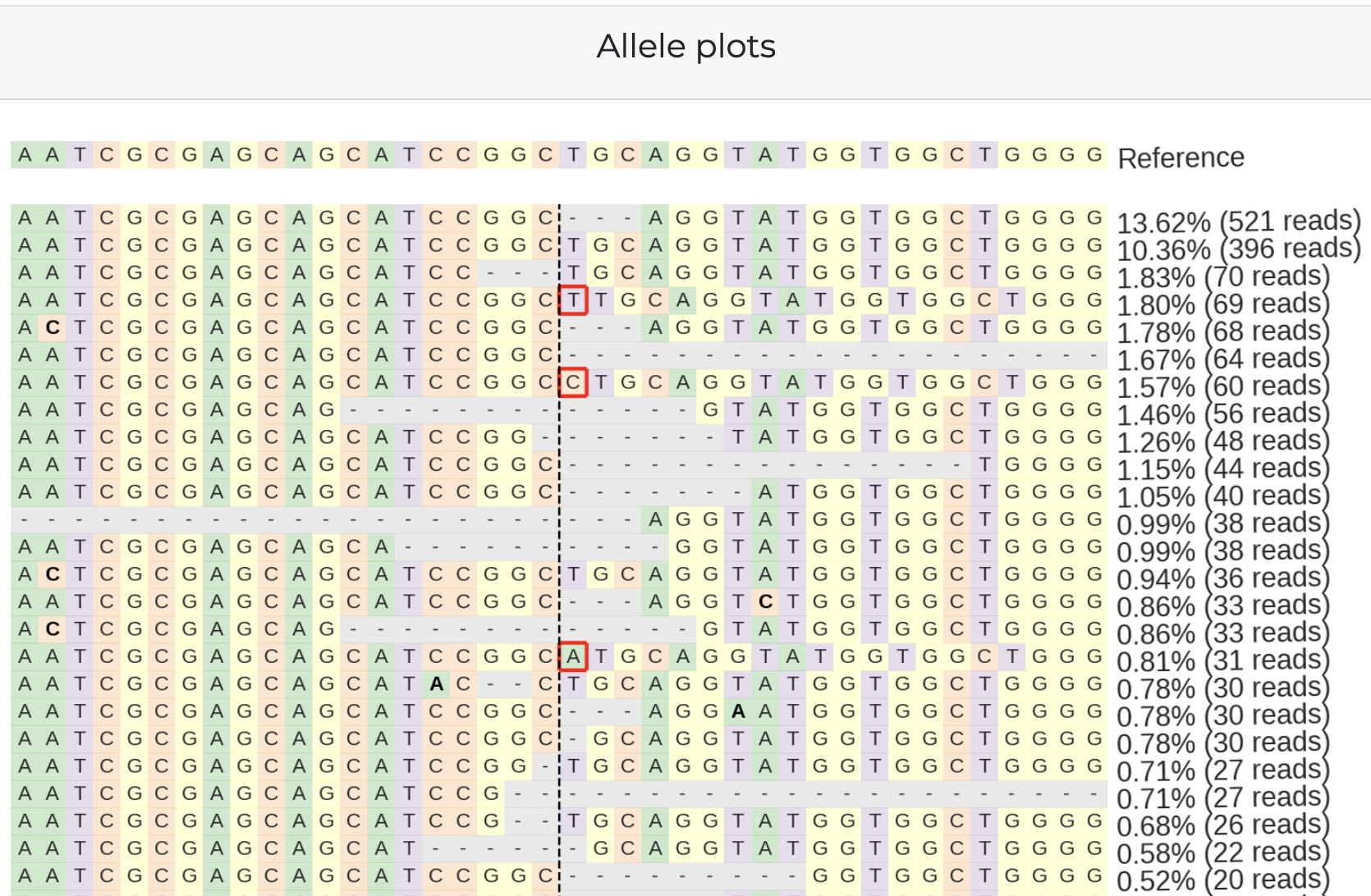

Figure S27: Crispresso report showing most frequent indels generated by NHEJ when targeting TSC2 exon 17. HAP1 cells were transfected with a gRNA targeting exon 17 of TSC2 cloned into a px458/9 family of vectors to co-express Cas9. Only about 10% of reads match the WT reference sequence. Most reads contain an indel suggesting highly efficient NHEJ.

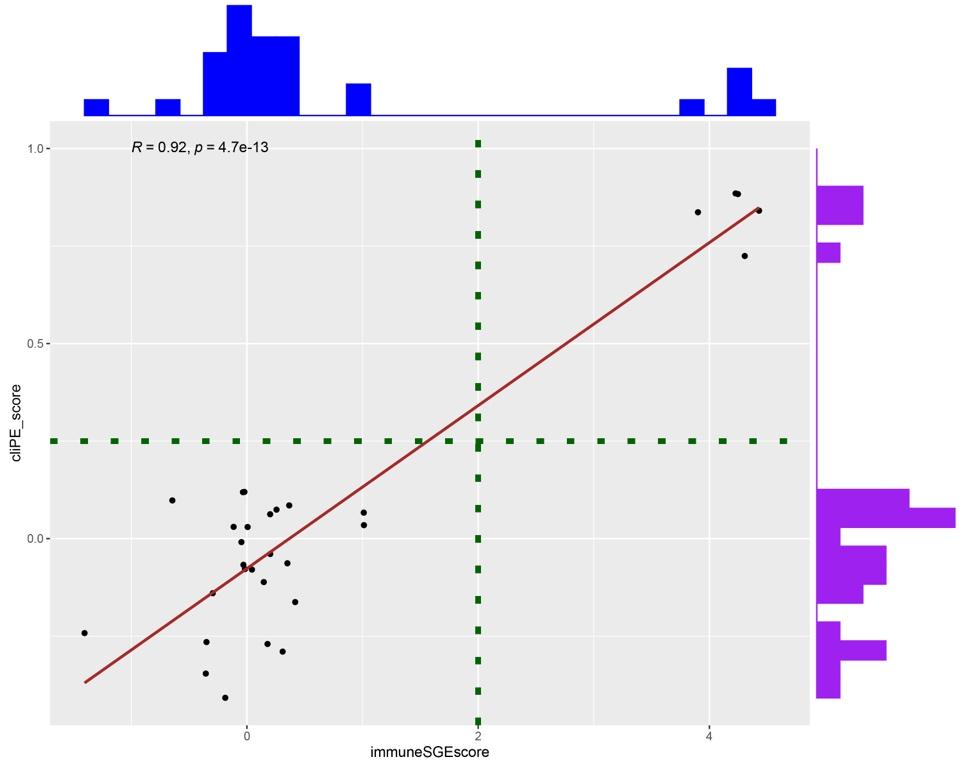

**Figure S28: MAVEs measuring TSC2 function are highly correlated, suggesting a lack of cell-type specific variant effects.**
